# Dynamic remodelling of cadherin contacts in embryonic mesenchymal cells during differential cell migration

**DOI:** 10.1101/2023.03.27.534409

**Authors:** David Rozema, Paul Lasko, François Fagotto

## Abstract

A fundamental aspect of morphogenesis is the capacity of cells to actively exchange neighbours while maintaining the overall cohesion of the tissue. These cell rearrangements require the dynamic remodelling of cadherin cell adhesions. Many studies have examined this process in tissues where it is driven by the joint action of cell protrusions and actomyosin contraction along the shrinking junction. However, cell rearrangements can also occur through differential migration. This mode of cell rearrangement, characteristic of mesenchymal tissues, is much less well understood. Here, we explore the prototypical case of the gastrulating Xenopus prechordal mesoderm, and provide the first detailed analysis at how cadherin contacts are remodelled and eventually disrupted in this type of tissue. Using a reductionist approach, including analysis of single contacts using a dual pipette aspiration setup, we unveil two concurrent mechanisms. Most cadherins are removed via “peeling”, i.e. disruption of the trans bonds and lateral diffusion out of the contact. In parallel, a remnant of cadherins concentrates at the shrinking contact, which is ultimately resolved by breakage of the link with the actin cytoskeleton, showing that the weakest link shifts at different stages of contact remodelling. Additionally, we observe recruitment of myosin peripheral to the shrinking contact, which influences the efficiency of the final detachment. Finally, manipulation of cortical tension indicates that the process is sensitive to the magnitude and orientation of the forces applied on the contact, revealing another key relationship between cell-cell adhesion and the cortical cytoskeleton. This study unravels a new modality of cell contact dynamics, which is likely to be widely relevant for highly migratory mesenchymal tissues.

## Introduction

Cell-cell rearrangements are critical for the morphogenesis of many tissues. The first major morphogenetic event in animal development is that of gastrulation where massive tissue-scale rearrangements lead to the repositioning of the three germ layers: ectoderm coating the exterior, endoderm occupying the interior, and mesoderm placed in between. Gastrulation in *Xenopus* is primarily driven by rearrangements at the cellular level that, in the case of the mesendoderm and prechordal mesoderm, proceed through a mesenchymal mode of migration where cells crawl using their neighbours as substrates (Huang and Winklbauer, 2018). This poses an interesting dilemma, as cell-cell adhesive contacts must be able to facilitate the overall cohesion of the embryo while still being amenable to remodelling to permit the cellular rearrangements that are required for gastrulation movements.

Classic cadherins are transmembrane glycoproteins and are the principal cell-cell adhesive molecules found across the animal kingdom (Hulpiau and van Roy, 2009). In this study, we focus on the type I classic cadherin, C-cadherin, the main cadherin expressed in the early *Xenopus* gastrula. Therefore, we will hereafter use ‘cadherin’ to refer to type I classic cadherins, although some of these concepts may be generalizable to all classic cadherins. Extracellularly, cadherins can form *trans* bonds with cadherins on opposing cell membranes and *cis* bonds with cadherins on the same cell both of which are important for developing mature adhesions (Brasch et al., 2012; Troyanovsky, 2022).

The cytoplasmic tail assembles the cadherin-catenin complex (CCC), which includes three adaptor proteins, β-catenin, α-catenin and p120 catenin. Cadherin binds to β-catenin, which binds to α-catenin, which in turn binds directly and indirectly to the actin cytoskeleton (Takeichi, 2014; Mège and Ishiyama, 2017), functionally linking the cytoskeletons of neighbouring cells. p120 catenin is involved in regulation of cadherin endocytosis (Takeichi, 2014; Katsuno-Kambe and Yap, 2020), and the whole CCC further interacts with various additional regulators that contribute to remodel the actin cytoskeleton at adhesion sites (Takeichi, 2014; Priya and Yap, 2015), as well as to regulate the strength of the cytoskeletal anchoring (Takeichi, 2014; Leckband and de Rooij, 2014; Mège and Ishiyama, 2017).

As an adhesion matures, cadherins organize into clusters, which is thought to strengthen adhesion (Brieher et al., 1996; Yap et al., 1998; Yap et al., 2015; Troyanovsky, 2022). Clustering is mediated extracellularly by the combined action of both *trans* and *cis* interactions (Wu et al., 2010; Wu et al., 2011; Wu et al., 2015), with additional intracellular contributions of p120 catenin (Ishiyama et al., 2010; Vu et al., 2021; Yap et al., 1998) and of the actin cytoskeleton (Truong Quang et al., 2013; Wu et al., 2015; Yap et al., 2015). Importantly, several of the interactions of the CCC are mechanosensitive (Buckley et al., 2014; Huang et al., 2017; Rakshit et al., 2012; Yao et al., 2014; Yonemura et al., 2010, Röper et al., 2018), and cadherin is enriched at the contact as a direct response to increased tension (Engl et al., 2014; Gao et al., 2018; Ladoux et al., 2010; Liu et al., 2010). It is clear that the contact can be reinforced in response to force, however, force is also required in order to facilitate cell rearrangements (Pinheiro and Bellaïche, 2018), leading again to the dilemma that mechanisms must exist to remodel and disrupt cadherin adhesions.

A large body of research exists that has attempted to identify the cellular and molecular engines that provide the force required to facilitate cell rearrangements. Two mechanisms were initially proposed. It was shown in epithelial models that oscillatory contractions of actomyosin along the remodelling junction were required (Bertet et al., 2004), while in mesenchymal models like the Xenopus chordamesoderm, cells exert traction on their neighbours with polarized protrusions (Shih and Keller, 1992). Though thought to be mutually exclusive, it was later demonstrated that both of these mechanisms are present and required in both epithelial and mesenchymal models (Huebner and Wallingford, 2018), with junctional actomyosin contraction observed in the Xenopus chordamesoderm (Shindo and Wallingford, 2014; Shindo et al., 2019; Weng et al., 2022), and basal protrusions observed in epithelial models (Williams et al., 2014; Sun et al., 2017). However, these mechanisms do not necessarily explain how cell rearrangements occur in all tissues. Another important mechanism is that of differential cell migration, where cells within a group migrate at different velocities than their neighbours, eventually outpacing them and applying tension on the contact that will stimulate remodelling (Huang and Winklbauer, 2018).

An additional avenue of research in cell rearrangements is how cadherin is removed from the cell contact. Several studies have suggested that this occurs via endocytosis (Levayer et al., 2011; Iyer et al., 2019; Katsuno-Kambe and Yap, 2020). Note that though endocytosis of cadherin is well established (Davis et al., 2003; Ireton et al., 2002; Le et al., 1999; Xiao et al., 2003) and has even been proposed to be directly responsible for disruption of the cadherin trans bond (de Beco et al., 2009; Troyanovsky et al., 2006), the issue remains unresolved, as removal of cadherin by endocytosis may be prevented when the cadherin is engaged extracellularly with other cadherins (in trans), and cytoplasmically with the cytoskeleton (Izumi et al., 2004; West and Harris, 2016). Alternatively, a different mechanism, termed ‘peeling’, has been proposed based on conceptual considerations (Evans, 1985; Garrivier et al., 2002). It is predicted that a tension applied tangential to the cell membrane acts primarily on the adhesive molecules at the periphery of the contact site. This would cause a gradual rupture of adhesion molecules, which would then free them to either be internalized or diffuse laterally on the free membrane. This model is consistent with recent evidence in Drosophila epithelia that forces applied perpendicular to the contact increase levels of cadherin at the contact, while forces applied parallel to the contact (shearing forces) decrease cadherin levels and thus ostensibly favour contact remodelling (Kale et al., 2018).

In this work, we sought to understand how cadherin is removed from remodelling cell contacts in the Xenopus prechordal mesoderm, a tissue that relies on differential cell migration to stimulate cell rearrangements (Evren et al., 2014). We probed several basic aspects of remodelling and disassembly of cadherin contacts. We used two reductionist systems, where dissociated cells were either studied while freely migrating on fibronectin (FN), or manipulated using a dual pipette aspiration (DPA) assay.

Previous studies using the DPA assay typically used it to measure the ‘separation force’ of cell-cell contacts (Daoudi et al., 2004; Chu et al., 2004; Chu et al., 2006; Maître et al., 2012; Slováková et al., 2022), which requires fast and large displacements to provoke instantaneous detachment of the contacts and assays equilibrium adhesion strength (Biro and Maître, 2015). As we wished to observe this force-sensitive remodelling on a physiological timescale, we used a different approach where the two cells were subjected to pulling forces over several minutes to match the speed of cell migration. We discover a stereotypical mode of cell contact remodelling involving two parallel processes: a removal of cadherin through peeling and subsequent diffusion on the free membrane, and a concomitant increase in density of the remaining cadherin at the contact. While peeling involved dissociation of the cadherin trans bond, the condensed residual cadherin appeared highly resilient, and final detachment was only resolved by abrupt rupture of the cytoplasmic link to the cytoskeleton, often involving the local recruitment of myosin. By altering cortical tension, we further uncovered how the two mechanisms of peeling and condensation depend on both the magnitude and orientation of the forces applied on the cell-cell contact.

## Results

### Remodelling of the cell-cell contact prior to separation involves both condensation and removal of cadherin

To visualize cadherin remodelling, we expressed fluorescently tagged C-cadherin by injection of the corresponding mRNA at the 2-cell stage. The tag was C-terminal, i.e. cytoplasmic. To keep total C-cadherin within physiological levels, we used our previously validated protocol, where the contribution of exogenous fluorescent cadherin was compensated by partial depletion of endogenous cadherin through co-injection of a morpholino antisense oligonucleotide (MO) (Canty et al, 2017). In this study we primarily used dissociated cells from induced mesoderm (IM), which was produced by the ectopic activation of the Wnt and TGFβ signalling pathways in ectodermal cells through expression of β−catenin and of a constitutively active activin receptor in the animal cap (Canty et al, 2017). IM cells closely resemble endogenous prechordal mesoderm in terms of adhesive and migratory properties and are more optically tractable due to their smaller size and reduced yolk content, making them an attractive model for this study (Canty et al., 2017).

Our initial approach involved using dissociated IM cells plated on FN and imaged using high speed 4D confocal microscopy. We mixed cells from two populations, one expressing C-cadherin-GFP and the other expressing C-cadherin-tdTomato, and imaged spontaneously forming doublets. Heterotypic doublets allowed us to unambiguously monitor cadherins from each of the two cells. We focused on single contacts formed by two cells that migrated in opposing directions, ostensibly applying a tension on the contact, which led to shrinking of the contact and eventual separation of the cells (Fig.1B; S1 video). Subsequent 3D segmentation (Fig.S1) allowed us to extract the area of the cell-cell contact and the total cadherin signal at the contact, which we then used to calculate cadherin density at each timepoint of the time lapse (Fig.1A’B,C). This quantification allowed to distinguish between two possible scenarios (Fig.1A,A’): first, cadherin molecules could be progressively removed as the contact shrinks, either through rupture of the extracellular trans bond and lateral diffusion along the membrane (a process we refer to as “peeling”), or through endocytic internalization, as proposed for contact remodelling of *Drosophila* epithelia (Levayer et al., 2011; Iyer et al., 2019). Either mechanism would result in a decrease in both contact area and total cadherin signal, and, if this removal was proportional to contact shrinkage, cadherin density at the contact may then remain constant (Fig.1A,A’). If, on the contrary, cadherins would not be removed from the shrinking contact, they would concentrate, which will be reflected by increased density as the contact shrinks, and, in the most extreme scenario, constant total signal (diagram Fig.1A,A’). The question would then be to determine how the increasingly dense contact would eventually resolve.

**Figure 1 –.**
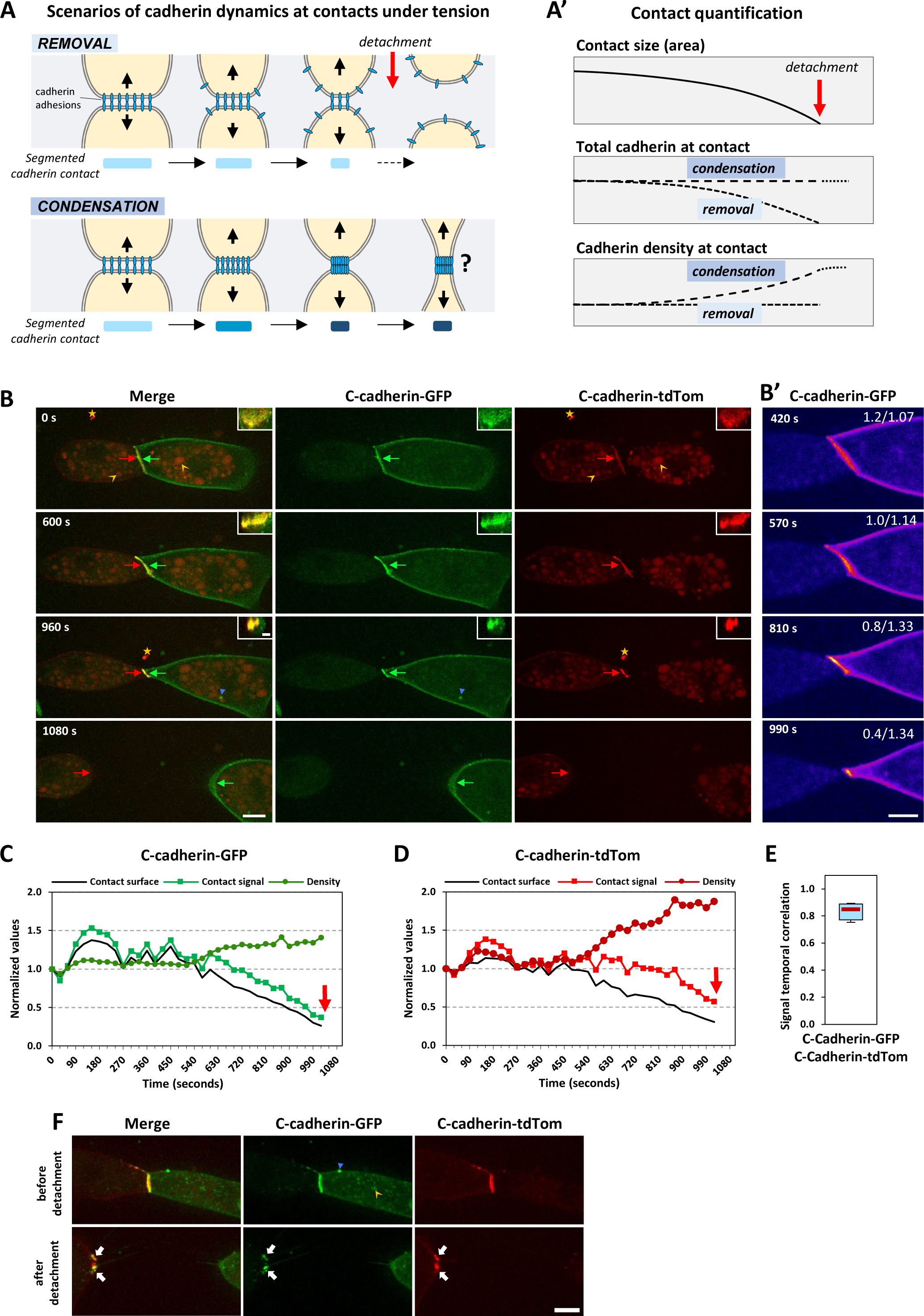
Contact remodelling during cell migration on FN shows a concomitant removal and condensation of cadherin from the cell contact. **(A)** Hypothetical opposite scenarios of cadherin dynamics in response to contact surfaces being pulled apart under tension. The blue ovals symbolize cadherin-based structures, abstracting multimeric organization, scale and actual membrane curvatures. The rectangles below each diagram represent the segmented contact with the cadherin density symbolized by shades of blue, with corresponding graphs of changes over time presented in (A’). Cadherin behaviour may fall between two extremes: Cadherin engaged in cell-cell adhesion may be progressively removed as the contact shrinks, either through peeling as depicted, or via endocytosis, not shown, such that its density at the contact remains unchanged, until the two cells detach (red arrow). Alternatively, cadherin may fail altogether to disengage, accumulating in an increasingly condensed structure. The question would then be how this dense and presumably very strong adhesive structure could be disrupted. **(B)** Contact shrinkage and detachment of two mesoderm cells plated on FN. Maximum intensity projections from a time-lapse. The cell on the right is expressing C-cadherin-GFP and the cell on the left C-cadherin-tdTomato. A z-stack of 12 μm was acquired every 30 seconds. The top right corner is a maximal projection of the YZ orthogonal view of the contact. Scale bar is 15 μm for Z projection and 5 μm for the YZ orthogonal view. Red and green arrows point at respectively C-cadherin-tdTomato and C-cadherin-GFP at the contact, and at the edge of the two cells after detachment. Concave orange arrowheads point to examples of the numerous autofluorescent cytoplasmic yolk platelets. Filled blue arrowhead: green cadherin intracellular spot. Orange star to free-floating red fluorescent debris. **(B’)** Sum intensity projection of sixteen z slices centred around the middle of the contact pseudo-coloured with a FIRE LUT to emphasize changes in intensity. Numbers indicate total relative contact cadherin/density. Examining the cytoplasm reveals no trace of intracellular cadherin in the vicinity of the contact. Scale bar, 5 μm. **(C,D)** Quantification of segmented contact surface, as well as total signal and average density for C-cadherin-GFP and C-cadherin-tdTomato of the cell double shown in (B). All values are normalized to the first timepoint. **(E)** Temporal correlation of the total cadherin signal comparing the slopes of C-cadherin-GFP and C-cadherin-tdTomato. Despite variations in magnitude of change, the overall patterns are highly consistent for both sides of the contact during remodelling. Comparison of seven doublets from five experiments. **(F)** Maximal intensity projection of a doublet prior and post detachment. White arrows highlight cadherin clusters containing cadherins from both cells after detachment of the cell contact, indicating that the contact ruptured cytoplasmically. Green and red speckles outside of the contacts come from signal at the cell surface. Scale bar 15 μm.

The analysis of 26 cell-cell detachments revealed that 20% of them occurred without detectable cadherin accumulation, thus with all cadherin being progressively removed as the contact was shrinking (example in supplemental Fig.S1D). The remaining 80% showed a hybrid process, with both a progressive loss of cadherins as well as a condensation (Figure 1B-D). The graphs show that though there was some fluctuation of the contact interface and of the corresponding total cadherin signal, the average density remained relatively stable over time (0-540s; Figure 1C,D). However, as the cells continued to migrate away from each other the contact started to shrink, and cadherin was removed from the contact (Fig.1B, YZ insets) with a concomitant condensation of the cadherin remaining at the shrinking contact (600s-1020s; Fig.1B-D). The use of differently labelled cadherins allowed an important observation: although the magnitude of the changes in fluorescence differed between the cells expressing the different tagged cadherins, the overall pattern was the same, which was demonstrated by the very strong correlation when comparing the slopes of contact signal between C-cadherin-GFP and C-cadherin-tdTomato (Fig.1E). Due to this strong correlation, we will only present the data for C-cadherin-GFP cells when heterotypic cell-cell contacts were imaged. Another feature frequently observed was the inefficient disassembly of the highly condensed remnant cadherin contact at later stages of contact remodelling (Fig.1F and supplemental Fig.S1E,F). This resulted in the stretching of long membrane protrusions between the cells, which would eventually snap, leaving cadherin clusters containing cadherin from both cells on one or both cell membranes. Since the cadherin constructs were tagged on the cytoplasmic tail, this implies that the final rupture of the adhesion did not occur at the extracellular cadherin trans bond, but between one of the cytoplasmic interactions. This phenomenon indicated that under some conditions, cadherins failed to disengage from dense clusters. Identical results were obtained for “homotypic” contacts established between two cells expressing the same tagged C-cadherin (GFP, supplemental Fig.S1D,E; tdTomato, supplemental Fig.S3).

These images showed that a large portion of cadherin was removed before the final detachment. This raised the question of the mode of cadherin removal. If cadherin was internalized, we would expect to see the apparition of cadherin positive endosomes in the cytoplasmic compartment, which was not the case. We did not detect any sign of endocytosis at the contact, despite the fast imaging. Occasionally, cadherin spots were observed, but the majority corresponded to pre-existing clusters that were seen diffusing on the plasma membrane (Fig.S1, orange arrowheads). At least part of them were likely remnants of disrupted contacts (see below). Few intracellular spots were found (Fig.1B, orange arrow), but could not be related to contact remodelling. It became evident that endocytosis could not account for the large and rapid cadherin removal in this system. We thus favoured the alternative mechanism, i.e. cadherin trans bond disassociation via peeling and lateral diffusion. However, the complex and ultrafast dynamics of the plasma membrane in the migrating cells prevented direct visualization of this process in these settings. Though this analysis of cells plated on FN provided key insights for our initial hypothesis, there were additional limitations to this approach. The detachment of the cells relies on random cell migration on FN. This inevitably leads to the application of inconsistent forces – in both magnitude, persistence, and orientation – on the cell-cell contact. Further, with cells constantly migrating, it was difficult to determine with certainty that we imaged the full ‘lifetime’ of a cell-cell contact i.e. starting from a tension-free equilibrium and moving towards increasing forces and contact disassembly.

### Separation of cells using dual pipette aspiration reveals a consistent detachment paradigm

To circumvent these issues, we opted for a more controlled approach, using a DPA setup, where two pipettes connected to a finely controlled negative pressure pump were used to apply a mild aspiration pressure on cells allowing for their precise manipulation (Biro and Maitre, 2015). Two dissociated cells were grabbed and brought into contact (Fig.2A). They were allowed to establish their adhesion for ∼5 minutes. This capacity to adhere extremely rapidly is characteristic of mesodermal cells (Rohani et al., 2014). After the cell-cell contact was established, doublets were imaged for several minutes at equilibrium, before displacing the cells incrementally away from each other until the contact was eventually ruptured. This approach had several benefits: Each doublet started at a similar level of adhesion, the application of force on the contact was consistent and controlled, and during the initial stages of imaging the cell-cell contact was ostensibly under no tension. The detachment behaviour observed with DPA was overall quite similar to that of cell doublets migrating on FN, with some nuances. During the initial stages of displacement, there was a period of relative stability in terms of contact interface, cadherin total signal average, and cadherin density (Fig.2B,C; S2 video; 0-180s). For cells plated on FN, this phase would correspond to the period where cells moved apart without yet signs of contact shrinkage (Fig.1C-E, 0-540s). This was followed by a period of gradually decreasing contact volume and intensity signal (Fig.2A,B, 180s-450s). During this period, cadherin density first raised mildly, then a characteristic sharp increase was observed, which in the example case of Fig.2A occurred after 15 μm of displacement (Fig.2B, 360s). After final detachment, condensates containing both red and green cadherins were found on one or both cells, consistent with cytoplasmic breakage of the adhesive structure (Fig.2B, 480s, white arrows, see below). By comparing the total cadherin signal at the contact and the average density between early and late stages of the remodelling process, we saw that in every case there was a decrease in contact signal and a parallel increase in average density (Fig.2D,E), supporting the hybrid model observed of cells on FN.

**Figure 2 –.**
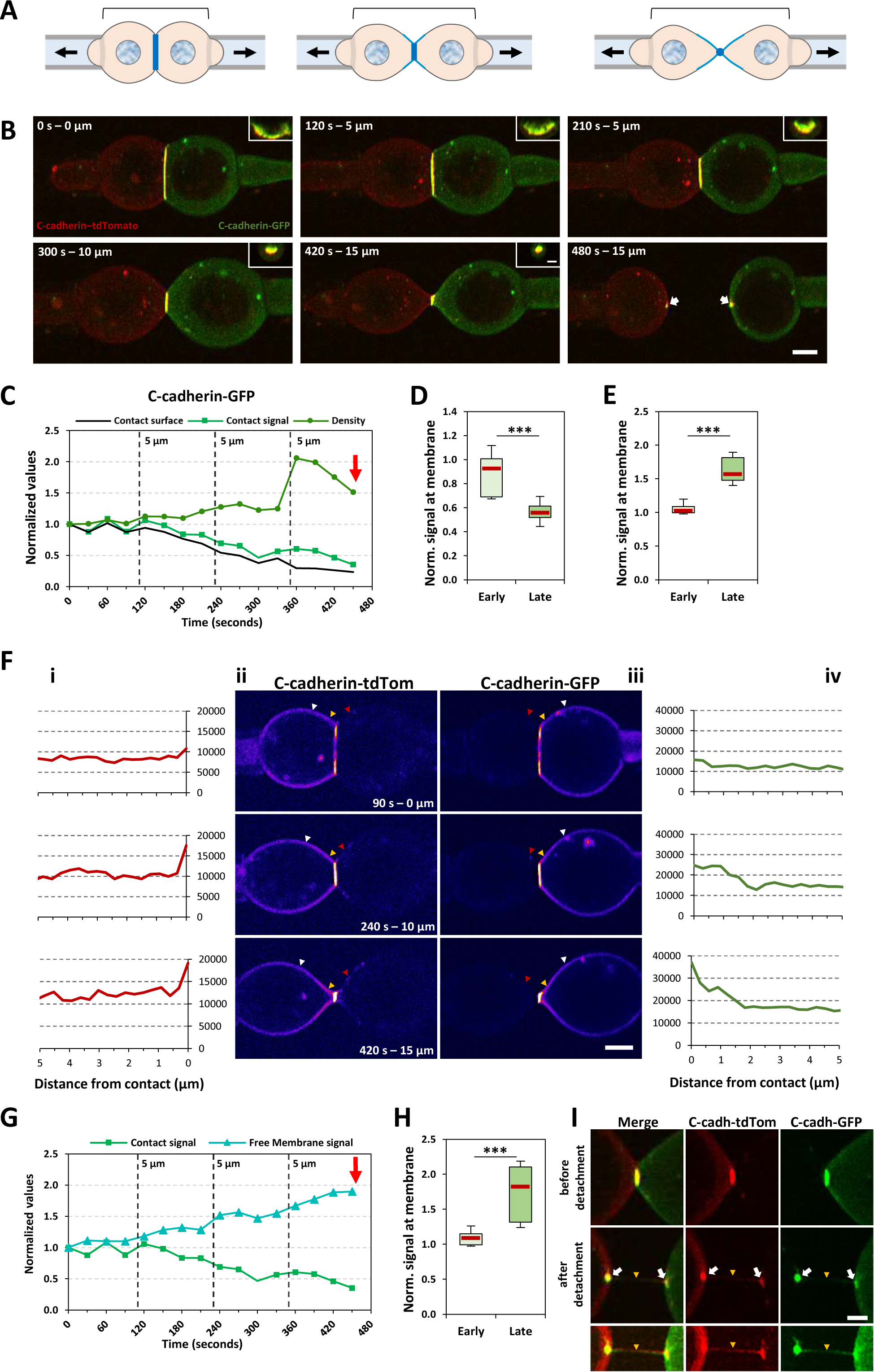
Displacement of cells using dual pipette aspiration reveals a consistent cell contact remodelling and detachment paradigm with cadherin removed from the contact diffusing on the free membrane. **(A) Diagram of DPA experiment.** The brackets indicate the stepwise displacement of the pipettes. **(B-E) General description of cadherin remodelling.** (B) Frames of a maximum intensity projection of a doublet being displaced using the DPA setup. The cell on the right is expressing C-cadherin-GFP and the cell on the left C-cadherin-tdTomato. A z-stack of 25 μm was acquired every 30 seconds. The time and displacement are noted in the top left for each frame. The inset in the top right is a max projection of the YZ orthogonal view of the contact. Scale bar is 15 μm for Z projection and 5 μm for the YZ orthogonal view. (C) Quantification of contact surface, total contact signal, and average density for C-cadherin-GFP and C-cadherin-tdTomato of the cell doublet shown in (B). All values are normalized to the first timepoint. Each vertical dashed line represents when the doublet was displaced by 5 μm. Red arrow marks detachment. (D,E) Quantification of the total signal at the contact and average density of cadherin comparing average values from two minutes (four timepoints) of the early stages of remodelling to the two minutes immediately prior to detachment. 9 doublets from 6 experiments. **(F-I) Quantification of diffusion of cadherin onto lateral membrane**. (F) (ii,iii) Sum intensity projections of seven Z slices centred around the middle of the cell contact, pseudocoloured with FIRE LUT to better visualize differences in intensity. Yellow arrowheads highlight increasing signal at the free membrane immediately adjacent to the contact, white arrowheads highlight the constant signal of the free membrane further from the contact suggesting increasing signal is coming from cadherin removed from the cell contact. Red arrowheads highlight either the lack of or very low levels of cadherin from one cell diffusing onto its neighbour, suggesting that the adhesive trans bond of cadherin is ruptured. Scale bar is 15 μm (i, iv) Line scans of the 5 μm adjacent to the contact that reveal increasing signal intensity at the free membrane. (G) Comparison of the normalized signal at the contact and at the adjacent free membrane of the doublet shown in (E), demonstrating that the membrane signal is enriched as the contact signal decreases. Vertical dashes indicate 5 μm displacements. Red arrow marks detachment. (H) Pooled quantification of the membrane signal of multiple doublets comparing the early and late stages of remodelling as described above. (I) Illustration of final cytoplasmic rupture. Max intensity projection of the frames immediately prior to and following detachment of a cell contact. White arrows indicate clusters that contain cadherin from both cells, demonstrating that the cadherin adhesive structure is ruptured cytoplasmically at the final step of detachment. The smaller bottom frames have adjusted brightness to emphasize presence of membrane tethers extending between the cells after detachment (yellow arrowhead). Scale bar is 5 μm. Statistical comparison for D, E, and H using a paired students t-test. Nine doublets compared from six separate experiments. P-values < 0.001.

An unexpected benefit of cell-cell separation using the DPA system was that it became clear how cadherin was being removed from the cell-cell contact. It was evident that as cadherin signal at the contact was decreasing, there was a large increase in signal along the free membrane (Fig.2F). The intensity of the membrane signal continued to increase throughout the later stages of the remodelling process (compare signal at white and orange arrowheads from 90s-420s; Fig.2Fii,iii; Fig.2G). Line scans spanning 5 μm of the membrane directly adjacent to the contact showed two characteristic, highly reproducible features: a steep slope adjacent to the contact, as well and a progressive increase over time of the base level of the signal in the next few microns (Fig.2Ei,iv). Comparing the early and late stages of multiple cell-cell detachments showed that the cadherin signal at the membrane increased in every case (Fig.2G), as the signal at the contact decreased (Fig.2D). Notably, the majority of cadherin was displaced back to the free membrane of the cell from which it originated (i.e. C-cadherin-GFP signal increased on the C-cadherin-GFP expressing cell, not the C-cadherin-tdTomato expressing cell), implying that the cadherin trans bonds were dissociating. Although occasional small clusters were found at the surface of the other cell (red arrowheads; Fig.2Eii,iii), this only occurred in a minority of cases (∼20%, discussed below). Consistent with experiments on FN, we did not detect any sign of cadherin internalization during the process.

As noted for cells on FN, we also observed that the final detachment, which generally involved a highly condensed cadherin contact, would often leave membrane tethers from each cell temporarily maintaining the connection between the cells (orange arrowhead, Fig.2I), and containing cadherin clusters positive for both colours of cadherin (white arrows; Fig.2I). Together these data clarify the observations from the doublets on FN and point towards a consistent mode of contact remodelling while under tension, involving a displacement of cadherin from the contact to the free membrane after peeling, while the cadherin remaining at the contact condenses before rupture of one of the cytoplasmic interactions of the CCC.

### The cadherin-catenin complex behaves as one unit during contact remodelling

We next turned our attention to the principal cytoplasmic binding partners of cadherin, the catenins. Using the same approach with the DPA system, we co-expressed C-cadherin-tdTomato with either p120 catenin-GFP or α-catenin-GFP. This allowed us to examine if other components of the CCC were removed from the contact prior to cadherin as it has been suggested for p120 catenin in *Drosophila* epithelium (Iyer et al., 2019), and α-catenin in zebrafish progenitor cells (Maître et al., 2012). If this was the case in our system, we would expect a decrease of signal of either catenin at least slightly preceding that of cadherin after displacement of the cells begins. Cadherin and p120 catenin co-localized at the cell-cell contact (Fig.3A,B; S3 video; merged images and YZ insets), and the remodelling events observed for cadherin, including decreasing signal at the contact and an increasing density (Fig.3A,D), were mirrored by p120 catenin (compare charts; Fig.3C). We found a strong temporal correlation between the slopes of cadherin and p120 catenin, well within the range of the correlation measured between the slopes of cadherin-GFP and cadherin-tdTomato (Fig.3I). Extending this analysis to α-catenin revealed similar results (Fig.3E,G,H; S4 video), again with strong correlation with remodelling of cadherin (Fig.3G,H,I). We verified that any cluster that would appear behind the contact (white asterisks; Fig.3A,E) was not intracellular, but rather located on the free plasma membrane on the lower planes of the image stack (see Fig.S1). After cells detached, the typical membrane tethers that were left between the two cells showed clusters containing both cadherin and p120 catenin (Fig.3B) or cadherin and α-catenin (Fig.3F). Combined with the presence of dual coloured cadherin clusters after detachment (Fig.2H), these observations were consistent with the final rupture occurring cytoplasmically, potentially at the α-catenin-actin link, or perhaps rupturing the actin cytoskeleton itself. These data suggest that the cadherin-catenin complex is remodelled as a whole, and that the catenins do not play specific roles to promote disassembly of the adhesion outside of the αcatenin/actin bond potentially being the weakest link at the final stage of contact detachment. Cytoplasmic rupture had previously been observed when separating cells by DPA in zebrafish, although in that case the CCC appeared to lose α-catenin (Maître et al., 2012). Levels of cadherin, β−catenin, and α-catenin have been previously found to remain proportional whether along cell free edges, clusters at cell contacts, as well as cluster-free tissue boundaries (Fagotto et al, 2013), indicating that at least in the early embryo of the frog, the CCC largely behaves as a single unit.

**Figure 3 –.**
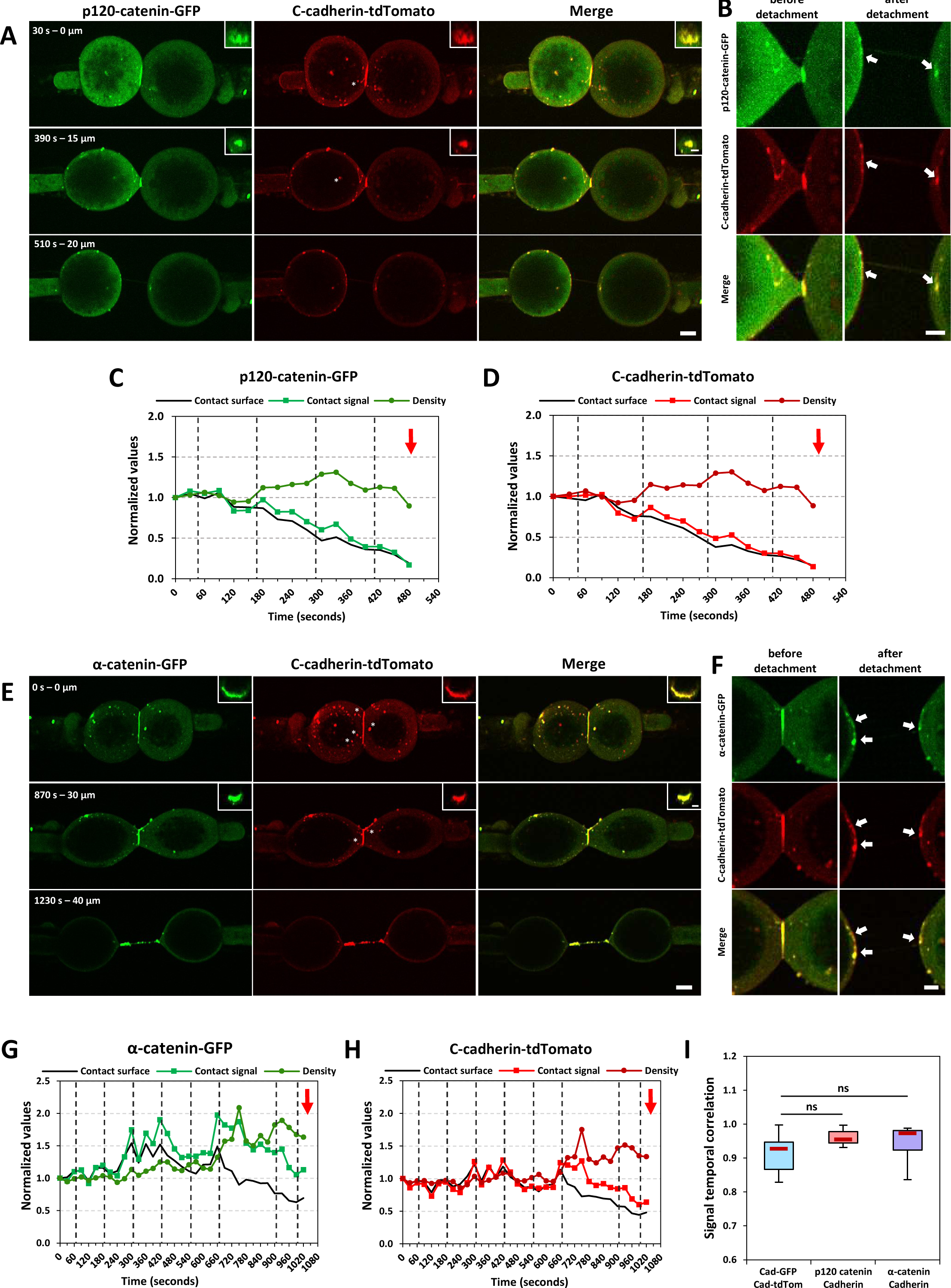
Imaging p120 and α-catenin demonstrates that the entire cadherin-catenin complex is simultaneously remodelled. (A, E) Frames of a maximum intensity projection of doublets being displaced using the DPA setup. Cells co-express C-cadherin-tdTomato with p120 catenin (A) or α-catenin (E). 25 μm z-stacks were acquired every 30 seconds. Time and displacement are noted in top left corner, while the top right is a maximum intensity projection of the YZ orthogonal view of the contact. Scale bars are 15 μm for Z projection and 5 μm for the YZ orthogonal view. (C,D,G,H) Quantification of contact surface, total contact signal, and average density for C-cadherin-tdTomato (D,H), p120 catenin-GFP (C), and α-catenin-GFP (G). Values are normalized to the first timepoint. Red arrow marks detachment. (B, F) Max intensity projection of the frames immediately prior to and following detachment of a cell contact. Arrows indicate clusters that contain cadherin and p120 catenin (B) or cadherin and α-catenin (F), further supporting the idea that the final detachment involves the rupture of one of the CCCs cytoplasmic interactions. White asterisks denote clusters that may appear cytoplasmic that are in fact on the free membrane on lower z planes of the image stack. See also Figure S3.1 (I) Temporal correlation of the slopes of the total contact signal comparing C-cadherin-tdTomato to either C-cadherin-GFP (n=9, 7 experiments), p120 catenin-GFP (n=10, 3 experiments), or α−catenin-GFP (n=10, 4 experiments). Statistical comparison using a one-way ANOVA with a post hoc Tukey test.

### Recruitment of myosin facilitates detachment of cells after condensation of cadherin

As actomyosin contraction stimulates cell-cell rearrangement and contact remodelling in other tissues, we were curious to explore its role in the prechordal mesoderm. We began by determining the spatial and temporal dynamics of myosin recruitment at the cortex of cells plated on FN, by co-expressing YFP-tagged myosin heavy chain IIA (MHCIIA-YFP) with C-cadherin-tdTomato. Interestingly, we often observed the appearance of a strong enrichment of MHCIIA during the terminal phase of detachment (Fig.4A,B; S5 video). Two important features characterized this recruitment of MHCIIA: 1) it occurred AFTER cadherin condensation had started, and 2) it was precisely located on the cortices immediately adjacent to the cell-cell contact, but never directly at the cadherin contact (YZ and XZ insets; Fig.4A), nor at the underlying cell-matrix interface (XZ insets; Fig.4A). Overall, ∼70% of imaged doublets recruited MHCIIA prior to the final detachment of the cells (see FigS3A,B for additional examples of detachments without and with MHCIIA recruitment, respectively). We could not extend this analysis of myosin in DPA settings, because MHCIIA-YFP expressing cells appeared to be very sensitive when kept in suspension and not provided with an adhesive substrate.

**Figure 4.**
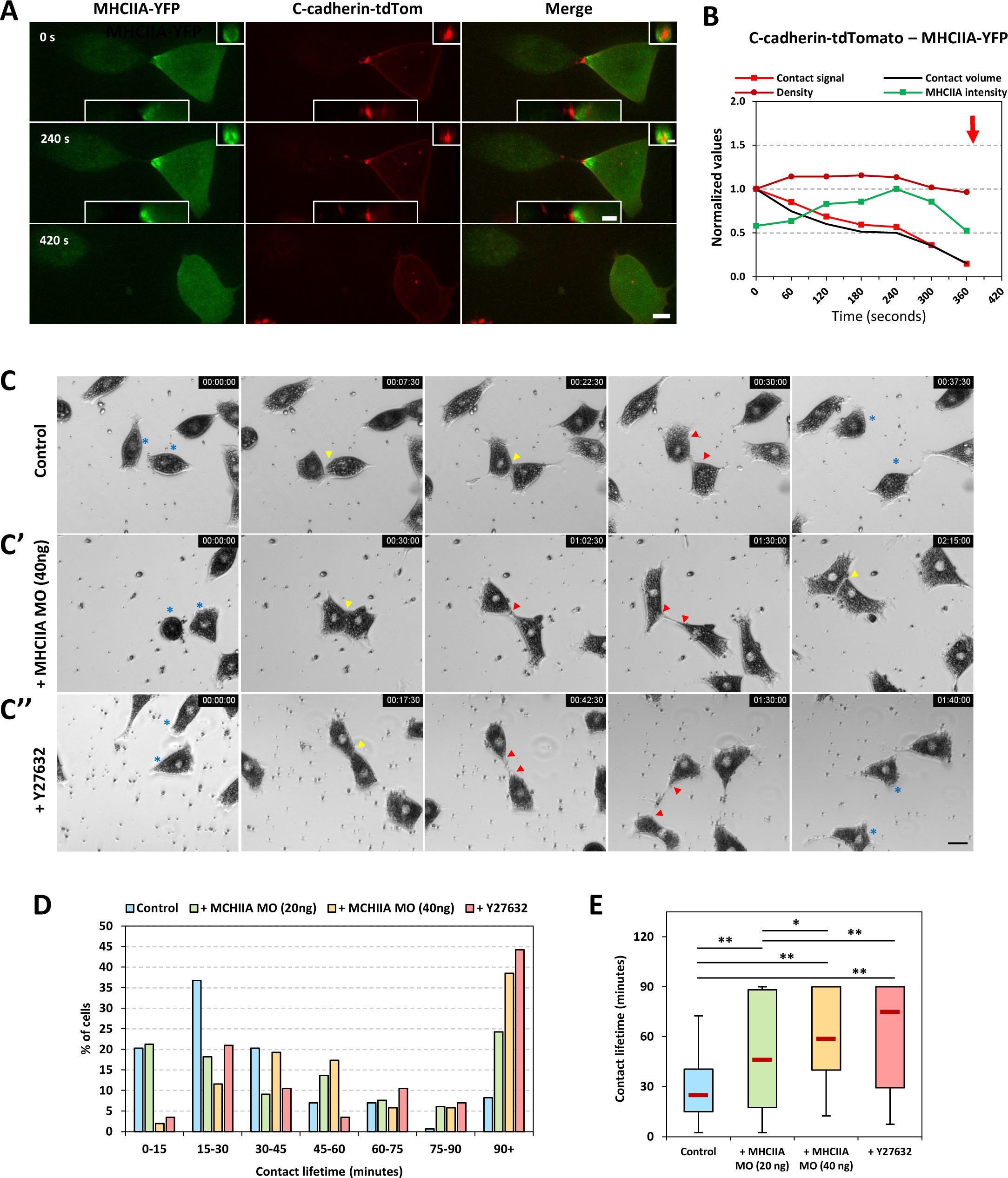
(A-B) MHCIIA recruitment adjacent to the cell contact after condensation of cadherin and prior to contact detachment. (A) Maximum intensity projection of a cell doublet plated on FN and co-expressing C-cadherin-tdTomato and MHCIIA-YFP, the right-hand cell having much higher expression. YZ and XZ orthogonal views are shown in the top right and bottom, respectively. A z-stack of 12 μm was acquired every 60 seconds. Scale bars are 15 μm for Z projection, 10 μm for XZ view, and 5 μm for YZ view. (B) Quantification of contact surface, total cadherin contact signal, and density, normalized to the first time point. MHCIIA-YFP signal is shown normalized to peak level. Red arrow marks detachment. **(C-E) Effect of myosin-mediated contractility on the ability of cells to detach from each other.** Contact lifetime assay: Dissociated cells were plated on FN and imaged every 2.5 minutes for a total of 150min. Newly formed contacts were identified, and their lifetime until cell-cell detachment was quantified. **(C)** Frames of representative time lapses. Time zero is the initiation of the cell contact and the final frame shows the cells just after detachment (or not in the case of C’). (C) Control cells; (C’) cells injected with 40 ng MHCIIA MO; (C”) cells treated with the ROCK inhibitor Y27632. Asterisks indicate the cells which formed the tracked cell contact, yellow arrowheads mark the contact, while red arrowheads mark the extended membrane protrusions that form between cells at late stages of contact remodelling prior to contact detachment. Scale bar, 30 μm. (D) Distribution of contact lifetimes in 15min bins for different groups. (E) Average lifetime for different treatment groups. Control (n=158, 4 experiments), MHCIIA MO 20 ng (n=66, 2 experiments), MHCIIA MO 40 ng (n=40, 2 experiments), Y27632 (n=86, 2 experiments). Statistical comparisons using a one-way ANOVA with a post hoc Tukey test. * p-value <0.05, ** p-value < 0.01.

Due to the consistent recruitment of MHCIIA to the rear of cells prior to detachment we asked whether myosin-mediated contractility was required for this process. To this end, we used a contact lifetime assay (Roycroft et al., 2018), where we plated dissociated cells on FN and measured how long two cells remained adhered to one another after their initial encounter. Control cells showed a rather stereotypical behaviour: once they encountered each other, they typically polarized and migrated away from each other, leading to shrinking of the cell-cell contact (black arrowheads; Fig.4C) followed by a brief phase where long protrusions bridged the two cells before finally detaching (Fig.4C, red arrowheads; S6 video). These residual connections corresponded to the membrane connections observed in our live confocal cadherin imaging (Fig.2H; Fig.S1E,F; S2B). Though there was a wide distribution of contact lifetimes (Fig.4D), this process generally lasted 20-40 minutes (Fig.4E). We then interrogated the role of myosin though morpholino (MO) knockdown (KD) of MHCIIA. Note that MHCIIB KD led to a strong inhibition of cell migration on FN, precluding to infer the impact on contact lifetime. However, MHCIIA MO, which did not significantly affect single cell migration (Fig.S2), drastically prolonged the persistence of the long membrane tethers (red arrowheads; Fig.4C’). This often completely prevented the detachment of the cells over the 130 min duration of the time lapses (Fig.4C’; S7 video). Increasing the levels of injected MO increased the frequency of cells failing to detach (Fig.4D), leading to increased average lifetimes at higher MO levels (Fig.4E). Treatment of cells with the ROCK inhibitor Y27632 also increased the persistence of the membrane tethers (red arrowheads; Fig.4C’’; S8 video), preventing cells from detaching (Fig.4D) and increasing average contact lifetime to a degree similar to injection of 40 ng of MHCIIA MO (Fig.4E). None of these treatments had any effect on the migration speed of single cells (Fig.S2). Together these data imply that, though detachment of cells can occur without detectable recruitment of MHCIIA (Fig.S3A), ROCK induced activation of MHCIIA contractility contributes to the final phase of detachment.

### Major features of contact remodelling observed in the prechordal mesoderm tissue explants

Intercellular migration involving frequent remodelling of cell-cell contacts can be readily observed in whole explants of dorsal mesoderm. While the complexity of such 3D system prevented the detailed analysis achieved with dissociated cells, major features that we had identified in minimalistic settings could be readily detected. In particular, quantification of cadherin levels and density at shrinking contacts showed the same profile as in cell doublets on FN or manipulated with DPA (Fig.5A-D; Fig.S4; S9, S10 video), including both a progressive loss of total cadherin signal, as well as an increased cadherin density, consistent with the hybrid mode of contact remodelling. Furthermore, we often observed myosin accumulation at the cortex peripheral to the condensed cadherin contact at the terminal stages of the remodelling process (Fig.5E.F). A similar recruitment of actin was recently observed in these same explants (Huang and Winklbauer, 2022), strengthening the evidence for the role of actomyosin contraction in contact remodelling in this tissue.

**Figure 5.**
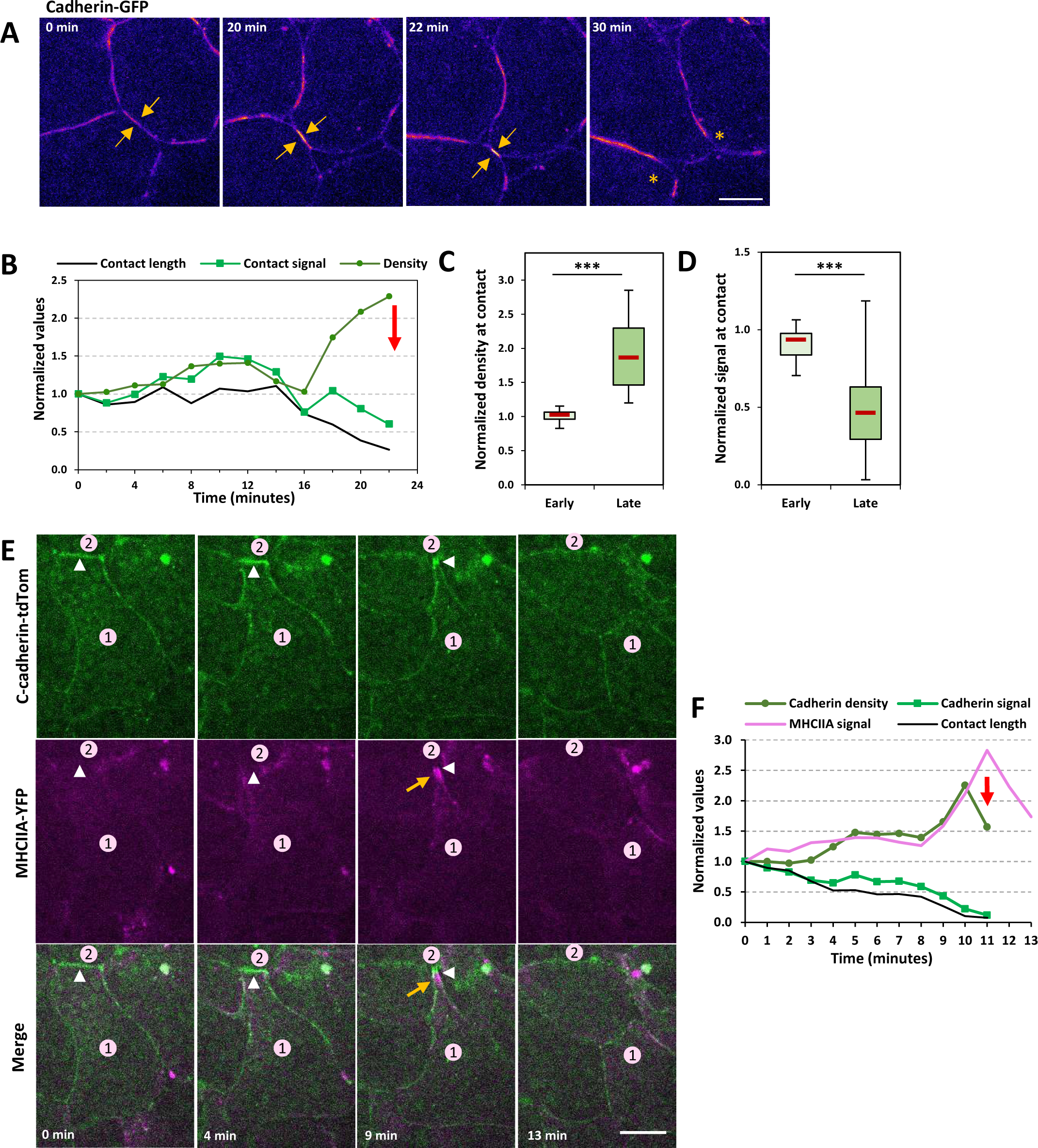
Cadherin and myosin at remodelling contacts during intercalation in whole mesoderm tissue. **(A-B)** Dorsal mesoderm explants expressing C-cadherin-GFP plated on FN. (A) Selected frames of time lapse movie. C-cadherin signal is presented using FIRE pseudocolours. Orange arrows point to the shrinking contact, asterisks mark the two cells after separation. Scale bar, 20 μm. (B) Corresponding profiles of the cadherin contact size, total cadherin and cadherin density. Red arrow marks detachment. (C,D) Normalized cadherin density and total cadherin at the beginning of contact shrinkage and at a late phase just before detachment. Average of 19 contacts, 3 experiments. **(E-F)** dorsal mesoderm explant coexpressing C-cadherin-tdTomato and MHCIIA-YFP. (E) Selected frames. Cell (1) in contact (white arrowhead) with its trailing neighbour (2) shows increased levels of MHCIIA at its rear (orange arrow at 9min) prior to detachment. Scale bar, 20 μm. (F) Profile of cadherin contact size, total cadherin signal and cadherin density, and MCHIIA-YFP total signal. Cadherin levels measured from line scans along the contact, while myosin levels measured by averaging the values from 10 μm line scans along both membranes peripheral to the contact. Red arrow marks detachment, MHCIIA-YFP signal persists briefly after detachment.

### Altering cortical tension impacts how cell contacts are remodelled

It is well established that cortical tension plays a central role in determining equilibrium adhesion strength (Maître and Heisenberg, 2013; Maître et al., 2012; Winklbauer, 2015; Slováková et al., 2022). Due to this interplay, we sought to explore if cortical tension influenced cell contact remodelling. Our first approach was to express a construct made of the actin-binding domain of utrophin fused with mCherry (UtrABD-Cherry). UtrABD is known to bundle actin (Belin et al., 2014) and therefore likely to increase cortical tension through enhancing the connectivity of the cortical actin network (Bendix et al., 2008; Ennomani et al., 2016). The aspiration pressures needed to stably hold the UtrABD-Cherry expressing cells were generally at least triple the pressure required to hold the control cells (∼250 Pa versus ∼80 Pa), which indicated that the cortex was much stiffer than that of the control cells. Upon displacing the UtrABD-expressing doublets the contact was only stable for a short time, then rapidly remodelled, leading to an abrupt loss of cadherin at the contact (120s; Fig.6A,B; S11 video). This mode of remodelling had three particularities. First, detachment occurred after a very short displacement (Fig.6H). Second, as the cell rapidly detached, there was little to no residual cadherin condensation, contrary to what was observed in control doublets (compare Fig.6A with Fig.2). However, a large number of small cadherin clusters formed on one or both of the cell surfaces prior to final detachment (blue arrowheads Y-and Z-projections; Fig.6A). These clusters were found over a large membrane area that roughly corresponded to the original extent of the cell-cell contact, suggesting multiple scattered sites of contact rupture. We call these events “early ruptures”, to distinguish them from the final rupture of condensed contacts described above. These early ruptured clusters were present in ∼70% of cases (Fig.6G), while the remaining ∼30% appeared to remodel purely due to peeling (Fig.6G). Although early ruptures were occasionally observed in control cells, they were in lower abundance, and loss of cadherin during the phase of contact shrinking could be largely attributed to peeling, concomitant to contact condensation (Fig.6G). Overall, UtrABD-injected cells required much smaller displacements to trigger detachment (Fig.6H), most cadherin was displaced before detachment, through cluster rupture and/or peeling, and the contact retained a stable cadherin density reflecting little to no condensation of cadherin (Fig.6B,I).

**Figure 6.**
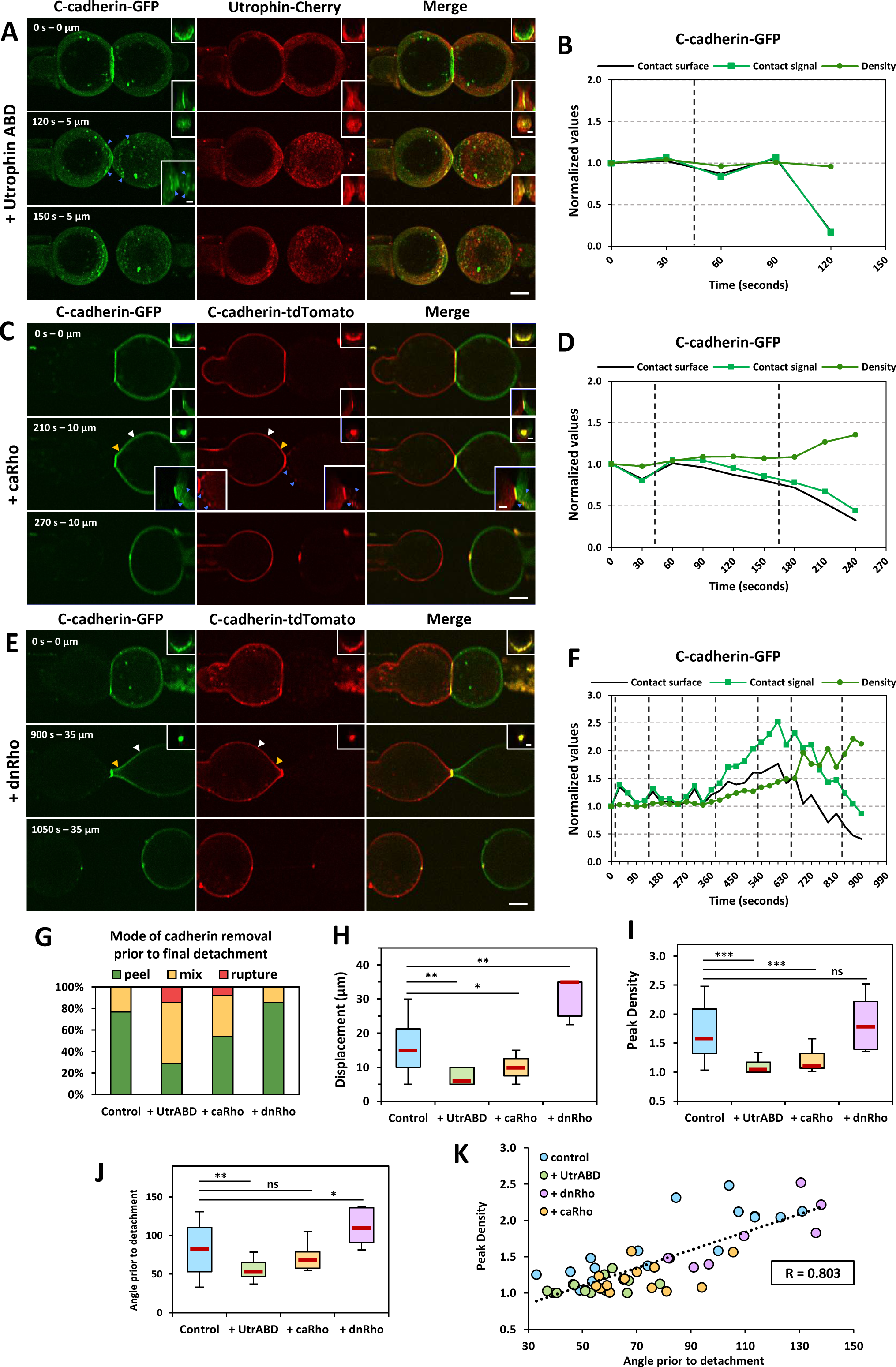
– Altering levels of cortical tension changes how cells remodel their contacts in response to force. (A) Max intensity projection of cell doublet co-expressing C-cadherin-GFP and utrophin-cherry. YZ and XZ orthogonal views are shown in top and bottom right corners, respectively. Blue arrowheads indicate clusters that ruptured cytoplasmically prior to the final detachment of the contact. (C,E) Sum intensity projections of cell doublets expressing C-cadherinGFP (left cells) or C-cadherin-tdTomato (right cells) and caRho (C) or dnRho (E). YZ and XZ (in C) orthogonal views are shown in top and bottom right corners, respectively. Blue arrowheads in (C) highlight ruptured clusters containing both coloured cadherins. Comparison of signal at the lateral membrane highlighted by yellow and white arrowheads show increasing signal on lateral membrane adjacent to cell contact. (A,C,E) 25 μm Z-stacks acquired every 30 seconds. Scale bar for Z-projections is 15 μm and 5 μm for the orthogonal views. (B,D,F) Quantification of contact surface, total cadherin signal and average density at the contact. All values normalized to the first time point. Vertical dashed lines represent 5 μm displacements. Red arrow marks detachment. (G) Characterization of the mode of remodelling during contact shrinking, showing the proportion of cells that remodelled via peeling, cytoplasmic rupture, or a mix of both. The final detachment, which in all cases involved cytoplasmic rupture, was not taken into account. (H, I, J) Quantification of displacement required for detachment, peak cadherin density prior to detachment, and angle between the cells prior to detachment. Control (n=17, 12 experiments), + utrophin (n=15, 6 experiments), + caRho (n=13, 5 experiments), + dnRho (n=7, 3 experiments). Statistical analysis using a one-way ANOVA with a post hoc Tukey test. * p-value <0.05, ** p-value < 0.01, *** p-value < 0.001. (K) Scatter plot showing the correlation between peak density and angle prior to detachment of the cell contact, all conditions combined. Pearson’s correlation coefficient: 0.803, n=52, p < 0.00001.

As an alternative way to increase cortical tension, we expressed a constitutively active RhoA (caRho) to stimulate ROCK-mediated myosin contractility. We noted a similar effect to that of UtrABD: smaller displacements were required before detachment of the cells occurred (Fig.6C,D,H), and little increase in cadherin density was observed (Fig.6D,I; S12 video). Analysis of dual colour cadherin doublets showed clusters from one cell appearing on the other cell prior to detachment, implying that the cytoplasmic interaction was broken when these early rupture events occurred (blue arrowheads; Fig.6C). Fig.6C also shows cadherin removal via peeling, visible by the cadherin gradient appearing next to the contact (compare orange and white arrowheads). Thus, contact shrinking appeared to involve early cluster rupture (Fig.S5B,C), peeling mechanisms, or a mixture of both. Scattered clusters were more frequent and more abundant in UtrABD-expressing doublets, indicating that their contacts were more likely to remodel through early rupture, while peeling was more widespread in the case of caRho-expressing cells (Fig.6G). Some cadherin condensation at the shrinking contact was observed (Fig.6D), but to a much lesser extent than in controls. Thus, both conditions of increased cortical tension favoured rapid disassembly of the contacts, with little to no cadherin condensation.

Taking the reverse approach, we expressed a dominant negative Rho variant (dnRho), and observed the opposite phenotype, as a larger displacement was required to detach the cells, not only compared to UtrABD and caRho conditions, but also to control cells (Fig.6E,F,H). The modality of early cadherin removal during contact shrinkage appeared to primarily rely on peeling (Fig.6G). Cadherin density significantly increased during contact remodelling, to levels equal or higher than in control cells, corresponding to a strong process of condensation (Fig.6F,I). It was recently demonstrated that increasing cortical tension can interfere with formation of a cell-cell contact, with higher tensions leading to smaller cell-cell contacts (Slováková et al., 2022). Measuring the relative contact size of all analysed doublets revealed that none of the perturbations used in our experiments affected contact formation (Fig S5D).

The tension applied to the cell contact in these experiments is a product of the displacement and the cortical tension. The increased rate of removal of cadherin from the contact and the smaller displacements required for detachment in the UtrABD and caRho injected cells could simply be due to the application of a larger tension. However, we also noted that in the stiffer cells, the angle of the vertex formed by the two cells at the contact was typically smaller than the angle in the control cells (Figs.6J; S5E), while cells injected with dnRho had more oblique angles prior to detachment. Together this implied that altering cortical tension alters the deformability of the cells (Fig.6J; S5E). Notably, when we plotted the peak cadherin density at the contact against the angle at the vertices immediately prior to detachment there was a strong correlation (R = 0.803, n = 52, p < 0.00001). Doublets that had acute angles prior to detachment often detached without any condensation of cadherin at the contact. On the other hand, doublets with oblique angles prior to detachment showed much higher levels of cadherin density (Fig.6K). These data demonstrate that in modulating the cortical tension of cells, we also alter how the cell contact is remodelled; increased cortical tension leads to rapid removal of cadherin through either accelerated peeling or cytoplasmic rupture, and lower cortical tension favours cadherin condensation and eventual removal by peeling. We emphasize again here that the final detachment in all cases still required rupture of the cytoplasmic bond. By changing the cortical tension of a cell, we do not only change the magnitude of the tension applied on the contact, but also the orientation at which it is applied, which also likely impacts the modality of contact remodelling.

Finally, after detachment of the doublets injected with UtrABD-cherry, we did occasionally see UtrABD-cherry colocalized with cadherin clusters suggesting that it could indeed be the cytoskeleton that is rupturing at the final phase of detachment (Fig. S5A). However, it was not as consistent or clear as the cadherin-cadherin or cadherin-catenin experiments. Additionally, as UtrABD affects the mechanical properties of the cells and their mode of contact remodelling, we cannot conclusively say whether this would also occur in control cells.

## Discussion

In this study, we describe the mechanisms involved in remodelling cadherin cell-cell contacts between mesoderm cells in response to tension. This process occurs naturally in the involuting prechordal mesoderm, where cells rearrange through intercalation driven by differential migration. We were able to unravel these mechanisms by using minimalistic settings, i.e. pairs of dissociated cells either freely migrating on FN, or subjected to controlled manipulation using a DPA system, which allowed us to dissect the dynamics of a single cell-cell contact in response to an applied tension. Using these approaches, we show that the system reacts to tension through two concurrent processes: cadherin disengagement from the contact through peeling or condensation of cadherin as the contact shrinks. Our data argue that, at physiological separation speed, peeling is a major route that removes cadherin, accounting for at least 50% of cadherin removal from the cell-cell contact. In parallel, cadherin condensation was also observed in most cases. It resulted in a shrunken contact densely packed with cadherin, which strongly resisted tension and was always resolved by an abrupt rupture of the cytoplasmic linkage. A typical detachment involved a hybrid mode combining both peeling and condensation followed by rupture.

Our observations of prechordal mesoderm contact remodelling depart from the mechanisms that have been described so far. Cadherin endocytosis is thought to play influence remodelling of adherens junctions of epithelia (Levayer et al., 2011; Hunter et al., 2015; Iyer et al., 2019), endothelia (Grimsley-Myers et al., 2020), collectively migrating astrocytes (Peglion et al., 2014), and during epiboly of the deep cell layer zebrafish (Song et al., 2013).While increased cadherin internalization has been suggested to specifically target the largest - and brightest - cadherin clusters (Truong Quang et al., 2013), we did not detect any sign of cadherin cluster translocation to the cytoplasm during contact shrinkage. Furthermore, while a recent study found that p120 catenin dissociated from cadherin in response to tension, leading to internalization of cadherin (Iyer et al., 2019), in our system, p120 catenin dynamics during contact disassembly were indistinguishable in time and space from that of cadherin, arguing that such mechanism does not play a significant role here. It seems that in *Xenopus* prechordal mesoderm, tension naturally exerted along the cell cortex by cells migrating away from each other (Evren et al., 2014, and our data Fig.1 and 5) coupled with contraction of the actomyosin cytoskeleton adjacent to the cell contact (this work; Roycroft et al., 2018; Huang and Winklbauer, 2022) provides an ample amount of force to disassemble a cadherin-based cell contact. As judged by the cadherin signal accumulating contiguous to the shrinking contact in our DPA experiments, peeling and lateral diffusion can amply account for the observed cadherin removal until final rupture. Although endocytosis does not appear to directly remodel cadherin adhesions here, it could potentially have a less direct impact through controlling the initial levels of cadherin present at the plasma membrane prior to contact formation (Davis et al., 2003; Ireton et al., 2002; Xiao et al., 2003).

A prior study using a DPA setup with zebrafish progenitor cells also reported the rupture of the cytoplasmic linkage when cells were rapidly separated (20µm/s; Maître et al., 2012). In that case, the α-catenin signal appeared to decrease before rupture, suggesting that it was the weakest point of the linkage. We did not observe any sign of release of any of the CCC components at any stage of contact remodelling, neither during the phase of peeling nor during final cytoplasmic rupture. We conclude that in our system, the weak point was more distal, either between the CCC and the actin cytoskeleton, or possibly within the cytoskeleton itself. Conceivably, the weakest link in the CCC at the final stage of contact remodelling may be context dependent, including cell types, but also kinetics of the detachment, which may impact the properties of the various molecular interactions. Our DPA pulling protocol was set to match physiological scales, i.e. the speed of normal mesoderm cell migration (∼2-3μm/min, suppl. Fig.S2, Kashkooli et al, 2021), and kinetics of cell-cell detachments observed for free migrating mesoderm cells, as well as for cells within the intact tissue (Figs.1,4,5; Evren et al., 2014; Huang and Winklbauer, 2022), providing confidence for the relevance of our results. Note that this mode of detachment by rupture is reminiscent of one mode of focal adhesion remodelling where after the final rupture paxillin remains at the substrate, implying that the extracellular integrin adhesive bond is not ruptured but rather one of the cytoplasmic interactions (Selhuber-Unkel et al., 2010).

Manipulation of cortical stiffness in our DPA experiments highlighted the importance of this parameter on the modality of contact disassembly. In this system, as the cells are displaced from each other, a tension is transmitted through the cortex onto the contact. The magnitude of this force is dependent on the cortical tension and on the size of the displacement. In other words, for the same displacement, a stiffer cell will apply a larger force on the contact than a softer cell. Doublets with high cortical tension rapidly detached with minimal displacement and condensation of cadherin was prevented, while doublets with low cortical tension required large displacements and involved strong condensation, leading to much higher cadherin density. Two parameters may explain these differences. Firstly, for stiffer cells, a larger force was being applied at a higher rate, which could impact the various molecular interactions of the CCC differently than the lower forces applied more slowly in softer cells. However, it was also clear that the softer cells were more deformable than the stiffer cells, as the angles between the cells at the contact prior to detachment was higher (Fig.6J,K; Fig.S5E). This is consistent with Rho-mediated contractility and actin organization being critical for setting resistance to deformation (Haase and Pelling, 2013). Further, the planar cell polarity protein, Prickle1, was recently shown to upregulate F-actin at the cortex in *Xenopus* prechordal mesoderm, and that Pk1 MO decreases F-actin levels at the cortex leading to a decrease in cortical tension. Pk1 MO also lowered the occurrence of neighbour exchange in dorsal lip explants and the cells were very elongated and deformed, seemingly unable to detach from their neighbours (Huang and Winklbauer, 2022). This study combined with our observations in dnRho injected cells suggest that regulation of cortical tension is an important factor to facilitate contact remodelling and cell rearrangements. The strong correlation between the peak density of cadherin at the contact and angle prior to detachment (Fig.6K) indicates that cell deformability may have a strong impact contact remodelling.

The process of contact detachment can thus be best captured by a model that integrates cadherin properties, cortical tension, and contact geometry (Fig.7). The proposed model is coarse-grained, but considers that the large-scale behaviour of cadherin adhesive structures is strongly dependent on the molecular properties of their components, in particular that of cadherin trans bonds and that of the cytoplasmic links with the cortical cytoskeleton (Fig.7A). Cadherin homotypic bonds bear strong resistance to tension applied parallel to the cadherin-cadherin axis (Leckband and Prakasam, 2006), but are likely to be very sensitive to forces applied orthogonally (*R*_ǀǀcad_ >> *R*_ꓕcad_). Simulations have suggested that forces with as little as 10° of a tangential component to the contact can be up to ∼50 times more disruptive to adhesion than a tensile force fully aligned with the adhesive bond (Chang and Hammer, 1996), and the notion has been experimentally supported in *Drosophila* epithelia (Kale et al., 2018). As cells are being pulled apart, tension *F*, transmitted through the cortex, is applied to the contact vertex at an angle (Fig.7A,A”). This tension can either cause the breaking of cadherin-cadherin bonds under load, or force cadherin-cadherin pairs to condensate. The angle, which is itself dependent on the stiffness of the cortex (Fig.7A’), influences the relative rates of each concurrent process: a large orthogonal component *F*_ꓕ_ favours bond breaking, thus peeling, while a large *F*_ǀǀ_ leads to condensation. The relationship between geometry and cadherin remodelling, validated in our measurements (Fig.6K), readily explains why we found rapid peeling and very little condensation in stiff cells, while soft cells built up condensed contacts (Fig.7A”,B). We systematically observed cytoplasmic rupture of the final large, condensed cluster (Fig.7A”,B), which is consistent with the superior resistance to rupture of trans bonds. In *Xenopus* prechordal mesoderm cells and in zebrafish progenitor cells (Maître et al., 2012), the bond most sensitive to *F*_ǀǀ_ happens to be somewhere within the cytoplasm, either at the catenin-actin interface, or in the cytoskeleton itself (*R*_cyto_ << *R*_||cad_). Thus, three major modes of contact disassembly can be defined (Fig.7B): peeling, condensation and rupture, and a hybrid of both, which was most commonly observed in our system. Note that while increased cortical stiffness could in principle yield a mode purely based on peeling, an additional phenomenon of rapid cytoplasmic rupture of adhesive clusters was observed with stiff cortices (Fig.6A,G; 7A’; S5B). This likely results from a sudden increase in force that surpasses both *R*_cyto_ and *R*_ꓕcad_, leading to a competition between cytoplasmic rupture and peeling to disassemble the contact. The high resistance of trans bonds along the axial direction may never be called upon since maximal resistance reached upon high tension parallel to this axis will first hit the lower resistance of cytoplasmic structures.

**Figure 7 –.**
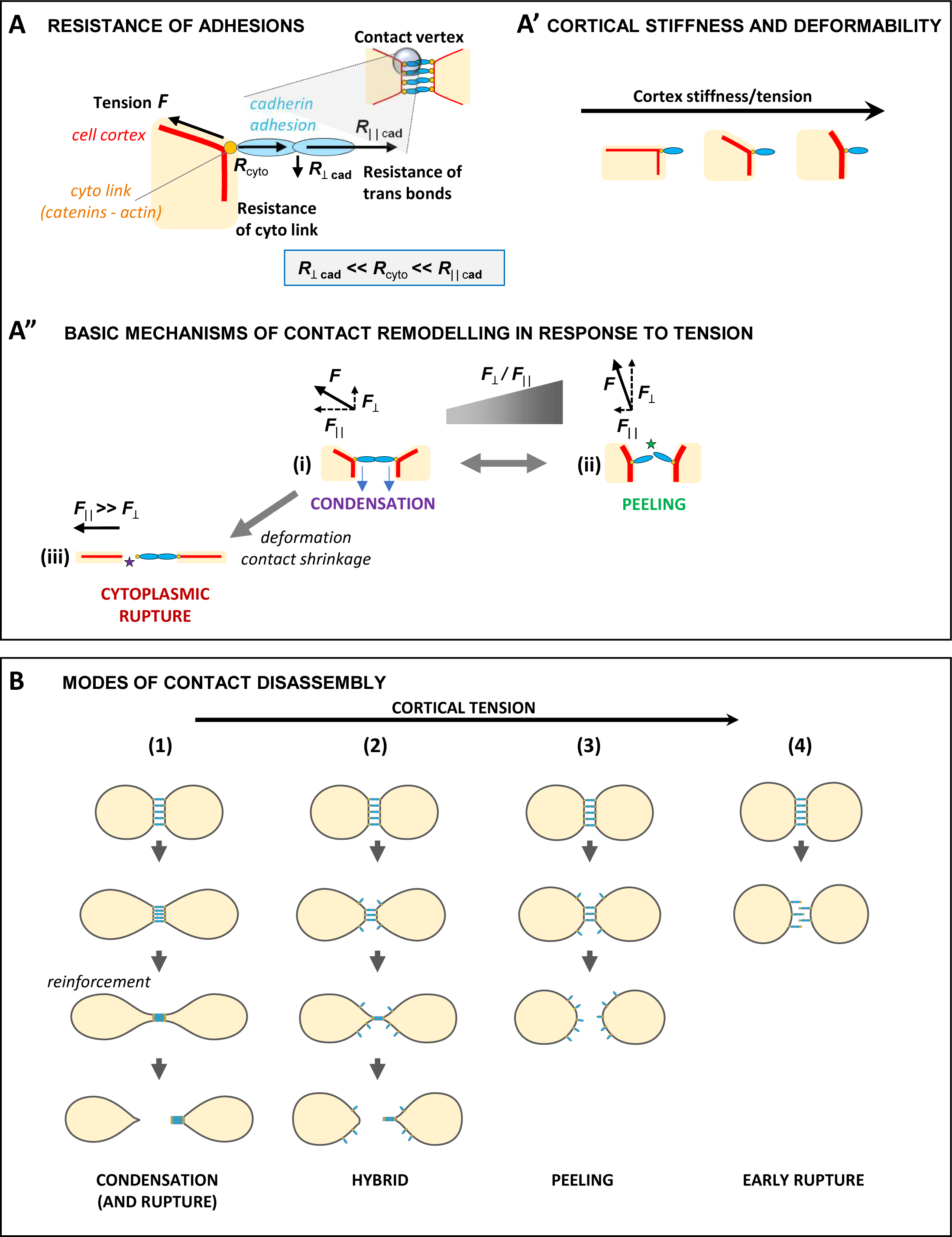
Model integrating the different modes of detachment and the impact of cortical tension. (A) Relationship between orientation of tensile force exerted at the edge of a contact and local resistance of cadherin adhesion. The cell cortex is represented by a red line. The cadherin adhesion structure (blue) is linked to cortex by the cadherin-catenin-based complex (orange). The plasma membrane, not depicted, is closely apposed to the cortex. Various components not to scale. For the sake of simplicity, relevant forces are represented without the corresponding force balance. The model considers that the resistance of cadherin adhesion to a tensile force *F* varies with geometry: It is highest parallel to trans bonds, lowest for a perpendicular force (*R*_‖‖cad_ >> *R*_ꓕcad_). *T*he resistance of the cytoplasmic link with the cortex (*R*_cyto_) lies intermediate between *R*_‖‖cad_ and *R*_ꓕcad_, thus *R*_‖‖cad_ >> *R*_cyto_ >> *R*_ꓕcad_. (A’) Inverse relationship between cortical stiffness and deformability. Cortical stiffness is symbolized by the thickness of the red line. For simplicity, no distinction is made between cortex at contact and at free edge, leaving aside regulation of cortical tension by cadherin adhesions at contacts. (A”) Basic mechanisms involved in contact remodelling. Pulling cell surfaces apart can lead either to peeling through breaking of cadherin trans bonds under load (ii) or shifting cadherin-cadherin pairs in the membranes, resulting in condensation (i). The relative rates of these two concurring processes are determined by orientation of the cortex tension (F_ꓕ_ / F_‖‖_). (iii) As final outcome of contact shrinkage and/or strong deformation, further facilitated by a weak cortex, *F* aligns with cadherins, facing maximal trans bonds resistance. As *R*_‖‖cad_ >> *R*_cyto_, the cytoplasmic link will rupture first. (B) Prototypical modes of cell-cell detachments, and their dependence on cortical tension. Four modes (1-4) are presented as schematic sequences from the initial situation before applying tension (top) to final detachment (bottom). (1) Cells with low cortical tension easily deform and most cadherin condenses. The resulting strong adhesive contact can only be resolved by cytoplasmic rupture, after extensive displacement. (2) Under normal conditions of intermediate cortical tension, a hybrid mode operates: Part of the cadherin pool peels and diffuses, while the rest condenses, and eventually breaks. (3) A stiffer cortex deforms less, favouring peeling, which on its own is sufficient for contact disassembly. (4) Increased cortex stiffness favours direct early cytoplasmic rupture upon short displacement. Experimental conditions in this study span through these four modes: Normal contacts were resolved mostly through the hybrid mode 2. As for DPA conditions (Fig.6), dnRho doublets behaved between modes 1 and 2, while caRho and utrophin doublets used variable contributions of modes 2, 3 and 4.

A striking aspect of contact dynamics of unmanipulated mesoderm cells is the wide variability: when examining the relationship between peak density and angle prior to detachment in the DPA experiments, the caRho and UtrABD injected cells occupy one end of the spectrum and dnRho the other, while the control cells span across the entire continuum. This variability likely reflects the known heterogeneity of cortical tension between individual cells (Canty et al., 2017; Kashkooli et al., 2021) and tissue stiffness between individual organisms (Zhou et al., 2015), and suggests that during gastrulation, mesoderm cells explore a wide spectrum of cortical tensions. It will be interesting in the future to test whether this heterogeneity is patterned and whether or not it facilitates cell-cell rearrangements during tissue morphogenesis. A related question to be addressed is the relative impact of peeling versus condensation-rupture on tissue dynamics. At first sight, peeling would appear as the most efficient mode of contact remodelling, and the requirement of cells to rupture the cytoplasmic link could be seen as an unwanted side effect of incomplete peeling. Yet one may wonder whether a given dose of condensation, imposing a tension threshold for cell detachment, may be a mechanism that contributes to set the right speed and coherence of the migrating tissue.

Such mechanisms will also depend on the force applied on the contact. Recruitment of myosin adjacent to the contact immediately prior to detachment suggests that the cells may occasionally require an extra force in addition to the tension exerted as cells move apart. Drawing an analogy to cell-matrix adhesion and migration is again informative as it is well established that cells migrating on an extracellular matrix can recruit myosin II to the rear of the cell to facilitate detachment from the substrate (Jay et al., 1995; Ridley et al., 2003), though other mechanisms can also be used (Cramer, 2013). However, myosin II-based contractility appears to be the dominant detachment force when larger forces are required, while other mechanisms are used when conditions are less stringent (Cramer, 2013). It is tempting to speculate that myosin II is used in a similar way at mesoderm cell-cell contacts, as it was consistently recruited after cadherin was already condensed at the contact and the forces required to detach are seemingly at their highest.

It is important to note that we have dissected contact remodelling in a well-defined topographic configuration, where the direction of migration is perpendicular to the orientation of the contact. The actual movements within whole tissues are obviously more complicated, with cells also crawling past each other laterally (migration parallel to the contact), and extensive radial cell intercalation initiated at gaps between cells (Huang and Winklbauer, 2018). Each configuration is likely to differ in how forces are applied on the contact and may produce different modes of contact remodelling. However, we believe that the mechanisms described reflect basic properties of cadherin contacts, and we predict that the principles discussed here will be widely applicable. Exploring these more complex situations and expanding the analysis to other embryonic tissues will be important to build a more complete view of tissue morphogenesis driven by differential cell migration.

## Materials and Methods

### Embryo preparation and injection

Plasmids and morpholino oligonucleotides (Genetools LLC) are listed in Tables S1 and S2 in the supplemental information section. mRNAs were synthesized according to manufacturer instructions (mMessage mMachine kit, Ambion). MOs and mRNAs were injected animally in the two blastomeres of 2-cell stage embryos for ectoderm targeting, or equatorially in the two dorsal blastomeres of 4-cell stage embryos for mesoderm targeting, at amounts listed in Tables S1 and S2.

### Mesoderm Induction

Embryos were injected animally at the two-cell stage with a mixture of mRNA including β-catenin (100 pg) and constitutively active Alk4 (1000 pg) as previously described (Canty et al., 2017).

### Chemical inhibitors

Y27632 was from Millipore. Stock solutions of were prepared in DMSO. They were used at a 1/1000 or higher dilution. Equivalent dilutions of DMSO were added to control conditions and had no detectable effect on cell and tissue properties.

### Microdissections and cell dissociation

All dissected explants for dissociation were taken either from the inner layer of the ectodermal animal cap after mesoderm induction. Keller explants were dissected from the anterior mesoderm at stage 10+ at the onset of involution. Dissections were performed in 1x MBSH (88mM NaCl, 1mM KCl, 2.4mM NaHCO_3_, 0.82mM MgSO_4_, 0.33mM Ca(NO_3_)_2_, 0.33mM CaCl_2_, 10mM Hepes and 10 μg/ml Streptomycin and Penicillin, pH 7.4. Single cells were dissociated in alkaline buffer (88mM NaCl, 1mM KCl and 10mM NaHCO_3_, pH = 9.5) (Rohani et al., 2014). All subsequent assays were performed in 0.5x MBSH buffer, at room temperature (23°C).

### Live microscopy

For cell migration and open faced Keller explants, dissociated cells or explants were plated on glass bottom dishes (Cellvis) that had been coated in for 45 minutes with 10 μg/mL bovine FN (Merck) followed by blocking with 5mg/mL bovine serum albumin (BSA). Cells and explants were then imaged on one of two spinning disk confocal microscopes: an Andor CSU-X1 with a iXon897 EMCCD camera controlled with Andor iQ3 software (Andor), or an Andor Dragonfly equipped with dual iXon888 EMCCD cameras and controlled by Fusion (Andor), both using a 40X 1.3 NA objective (Nikon).

### Image analysis and quantification

All confocal images were deconvolved using either Huygens deconvolution software (SVI) or Fusion (Andor). Cadherin, p120-, and α-catenin signal was segmented using the 3D imagine software Imaris (Oxford Instruments) to extract the total signal and volume at the cell contact.

The signal of cadherin at the free membrane was determined using sum intensity projections of the seven Z planes surrounding the centre of the cell contact. Line scans were drawn from the cell-pipette interface of one cell to the other, and 5 μm on either side of the contact was considered for analysis.

Temporal correlation was determined by comparing the slopes of the total signals of p120 catenin-GFP, α-catenin-GFP, or C-cadherin-GFP to that of C-cadherin-tdTomato at each timepoint and calculating the correlation coefficient.

### Dual pipette aspiration assay

Dissociated cells expressing fluorescent constructs were plated on a glass bottom custom-made chamber blocked for 45 minutes with BSA. The DPA assay was setup as described elsewhere (Biro and Maître, 2015), though instead of using large, fast displacements to rapidly detach cells and determine separation force, we incrementally displaced cell doublets 5 μm at a time while simultaneously imaging the double with confocal microscopy. We waited two to three minutes between each displacement to observe how the contact responded to increasing levels of force, and to imitate the speed at which cells migrate within the mesoderm or when freely migrating on FN (Kashkooli et al., 2021). Cells were manipulated with custom made pipettes with diameters from 13-17 μm with a 15° bend (Sutter Instruments) that were coated with BSA. Holding pressures of 80-400 Pa (condition dependent) were used to stably aspirate the cells for the course of the displacement protocol. Pressure was controlled using a Microfluidic Flow Control System and the Maesflow software (Fluigent), and the pipettes were manipulated using PatchStar Micromanipulators and the LinLab2 software (Scientifica). A 25 μm Z-stack was acquired every 30 seconds until the cells detached using the Andor Dragonly microscope described above.

### Contact lifetime assay

Dissociated cells were plated on FN coated glass bottom dishes and left to adhere for 30 minutes then imaged every 2.5 minutes for 150 minutes using a bright field inverted Olympus IX83 microscope equipped with a scMOS ZYLA 4.2 MP camera and a 10X 0.3 NA PH1 objective. Single cells that encountered another cell withing the first 40 minutes of observation were tracked and the duration of the cell-cell contact was measured. We counted any lifetime longer than 90 minutes as 90 minutes, as this was often near the point where cell viability decreased so any longer lifetimes could have been due to low cell viability. Inhibitors were added at the start of the timelapse, and equivalent concentrations of a DMSO vehicle control were added to the other conditions. When MHCIIA MO was injected the control cells were injected with an equivalent amount of control MO.

### Migration assay

Dissociated cells were plated on fibronectin-coated glass bottom dishes and left to adhere for 45-60min, then imaged every 2.5min for 100-170min using a bright field inverted Olympus IX83 microscope (10X UPFLN 0.3NA PH1 objective) and a scMOS ZYLA 4.2 MP camera. Chemical inhibitors were added after four frames (10min) after the beginning of the time lapse. Addition of the inhibitor was set as time zero. The path of individual cells that did not establish contacts with neighbouring cells were manually tracked using ImageJ software. Average speed corresponds to the average of the speeds calculated between each consecutive time point.

## Acknowledgements

We thank Rudi Winklbauer and Jean-Léon Maître for critical reading of the manuscript. We thank the Montpellier Rio Imaging platform for technical support. This work was funded by a Labex EpiGenMed Montpellier Chair of Excellence awarded to FF, (https://www.epigenmed.fr/index.php/funding/chair-of-excellence), and by grants awarded to FF from the Agence Nationale de la Recherche (ANR-14-ACHN-0004– ICM and ANR-21-CE13-0042-01, https://anr.fr/), as well as a grant MOP-130350 from the Canadian Institute of Health Research (https://cihr-irsc.gc.ca), awarded to PL and FF. The funders had no role in study design, data collection and analysis, decision to publish, or preparation of the manuscript.

## Supporting Information

**Supplemental Figure S1.**
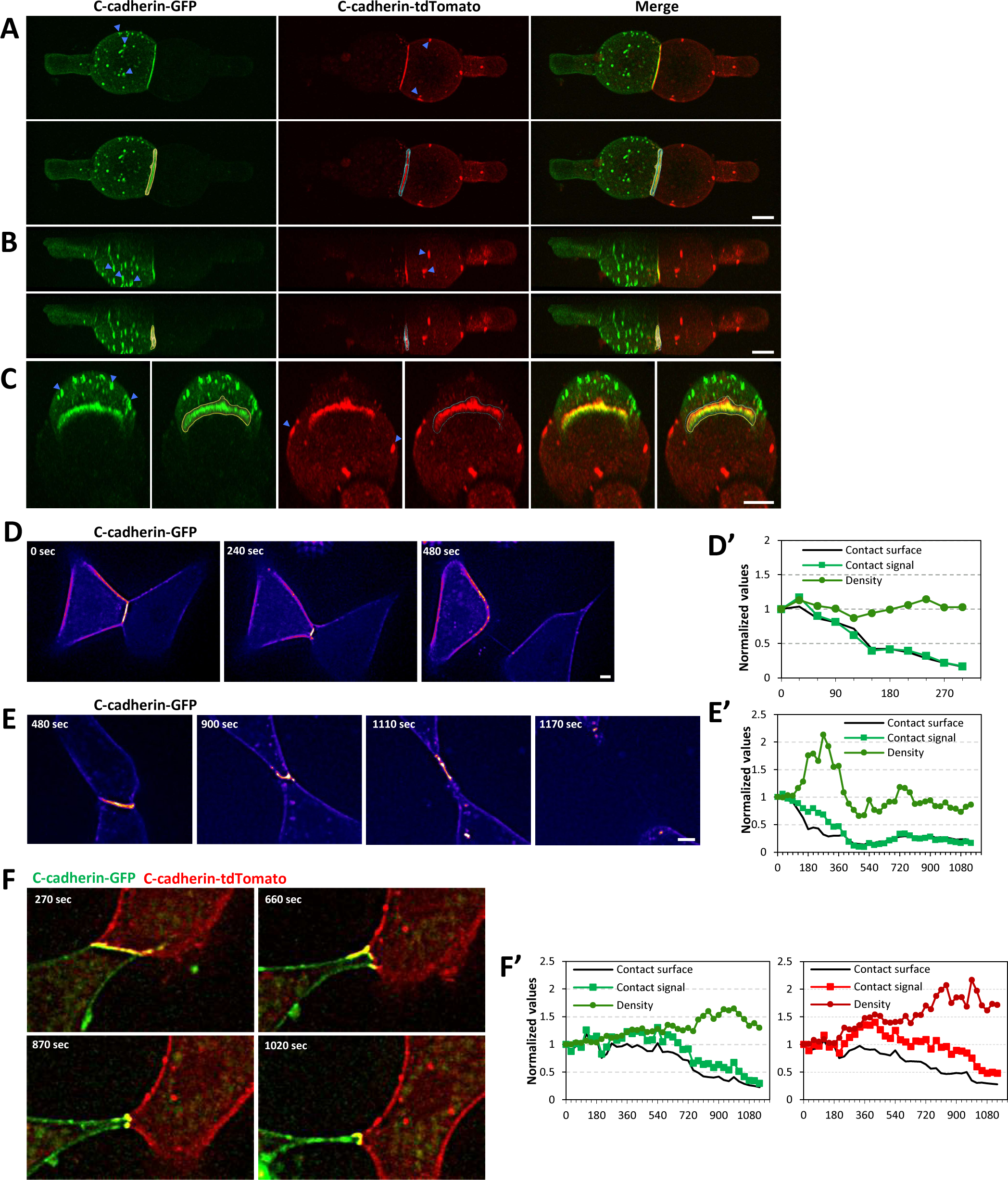
(related to Figure 1, 2, and 3). A) 3D segmentation of cadherin signal. Example of cells held using DPA assay visualized in the 3D imaging software, Imaris. (A) Top-down view of cells. Bottom frame shows the segmentation output, yellow outline for C-cadherin-GFP and blue outline for C-cadherin-tdTomato. (B) Side view to highlight height of the contact. (C) View at angle to emphasize the shape of overall shape of the contact. Frames on right show segmentation output. Blue arrowheads highlight clusters that appear cytoplasmic from topdown or side views but are in fact clearly localized on the free membrane from the angled view. D-F) Example of detachments of cell doublets on FN. D) Detachment mostly through peeling. Pair of cells both expressing C-cadherin-GFP, intensity shown as FIRE pseudocolours. D’) Quantification of cadherin at the contact shows a constant decrease in total cadherin signal without noticeable change in cadherin density. E) Hybrid detachment ending with a long phase of stretching of the residual contact. E’) Quantification shows parallel decrease in total contact signal and large increase in density, with a long trailing phase before detachment. F) Example of hybrid detachment for a pair of cells expressing C-cadherin-GFP and C-cadherin-tdTomato. Density steadily increases while total signal is only slowly decreased as the contact region stretches. Scale bars, 10 μm.

**Supplemental Figure S2.**
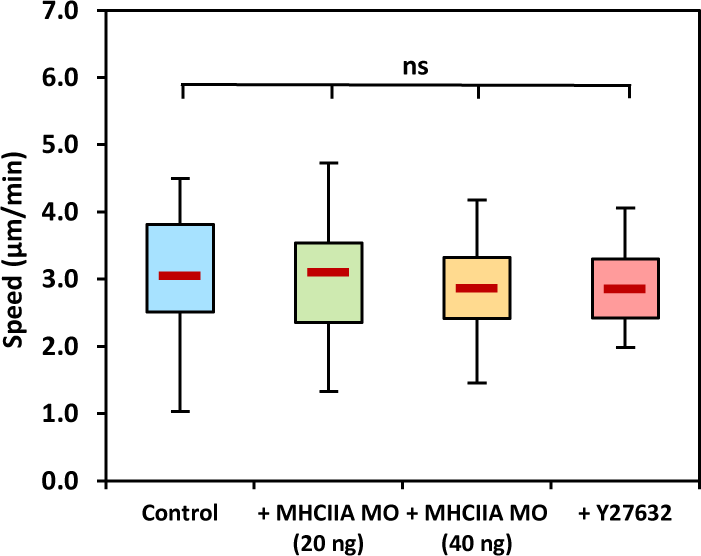
(related to Figure 4) Migration speeds of single cells plated on FN. Migration speeds of control, +MHCIIA MO (20ng), +MHCIIA MO (40ng), and +Y27632 treated single cells. No treatment had any effect on the rate of migration. Control: n=50(4); MHCIIA (20ng): n=39(2); MHCIIA (40ng): 35(2); Y27632: n5=36(2). # of cells (# of experiments). Statistical analysis using a one-way ANOVA with a post hoc Tukey test.

**Supplemental Figure S3.**
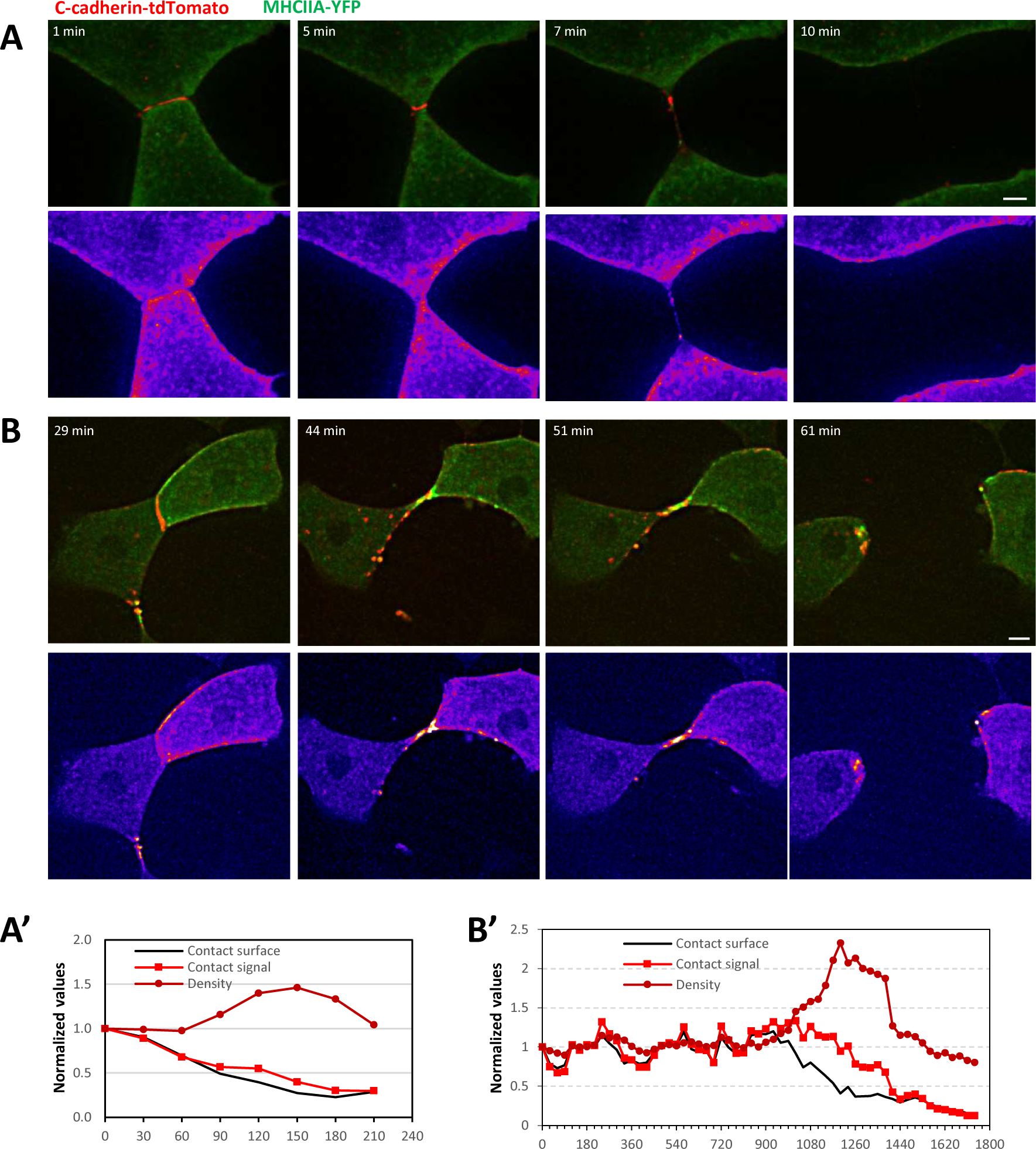
(related to Figure 5). (A) Example of detachment without noticeable myosin recruitment. The contact of these two cells expressing C-cadherin-tdTomato and MHCIIA-YFP detaches in a hybrid mode. (B) Example of detachment involving a long phase of stretching, correlated with strong recruitment of MHCIIA. The lower panels show the myosin signal pseudocoloured using FIRE LUT. (A’ and B’) Quantification of C-cadherin-tdTomato. Scale bars, 10μm.

**Supplemental Figure S4.**
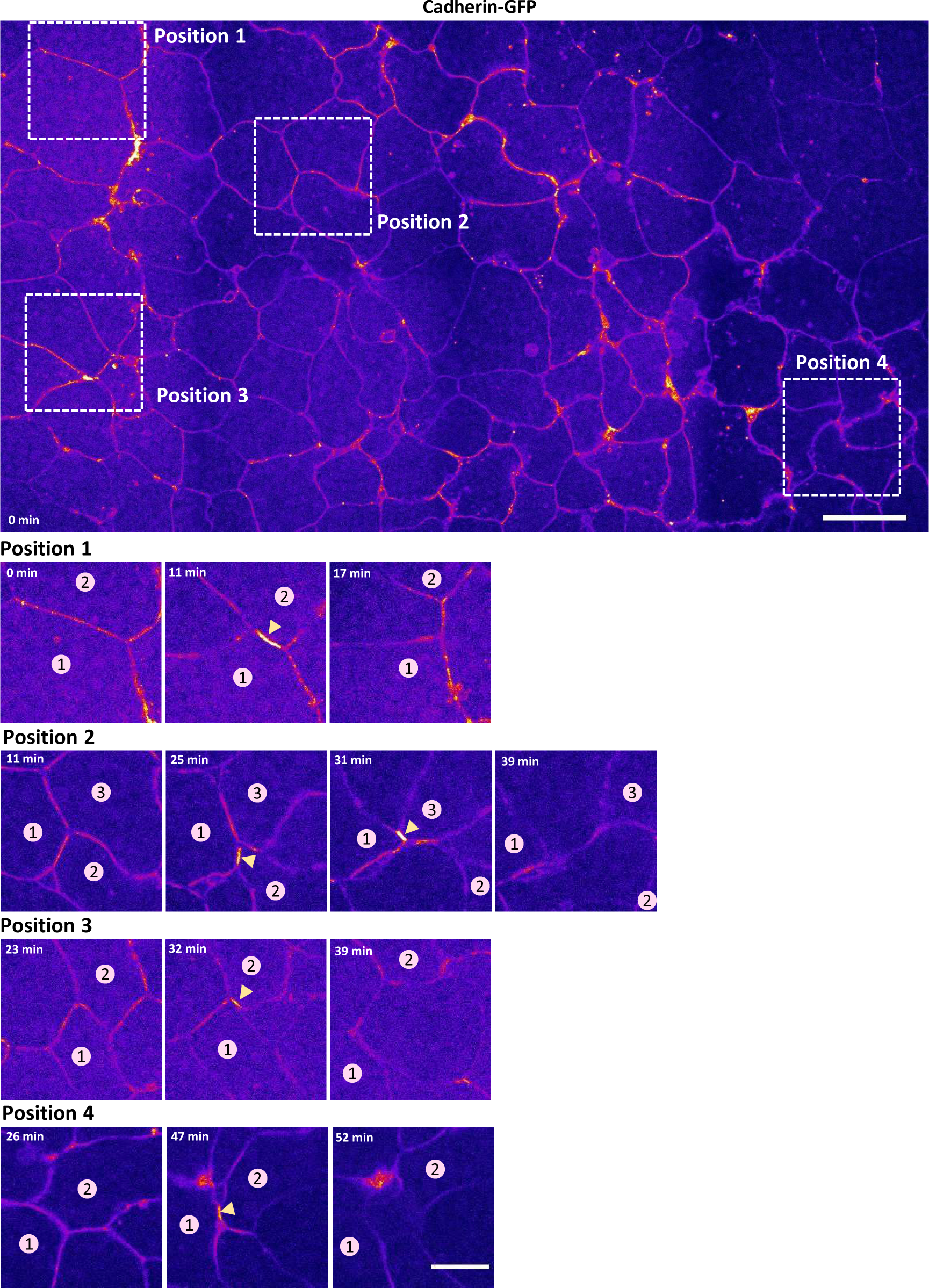
(related to Figure 5). Examples of intercalations with transient cadherin condensation just before contact detachment. Large field of view from a time-lapse of an open face mesoderm explant (OFE) expressing cadherin-GFP, shown as FIRE pseudocolours. 4 positions were selected to illustrate typical intercalation events, shown in detail in 3-4 time frames. The numbers mark the two cells initially in contact. In position 2, two consecutive intercalations are seen, first between cells 1 and 2, then between cells 1 and 3. Yellow arrowheads point to the condensed cadherin contact. Scale bars: Large frame 40 μm, small frames 20 μm.

**Figure S5.**
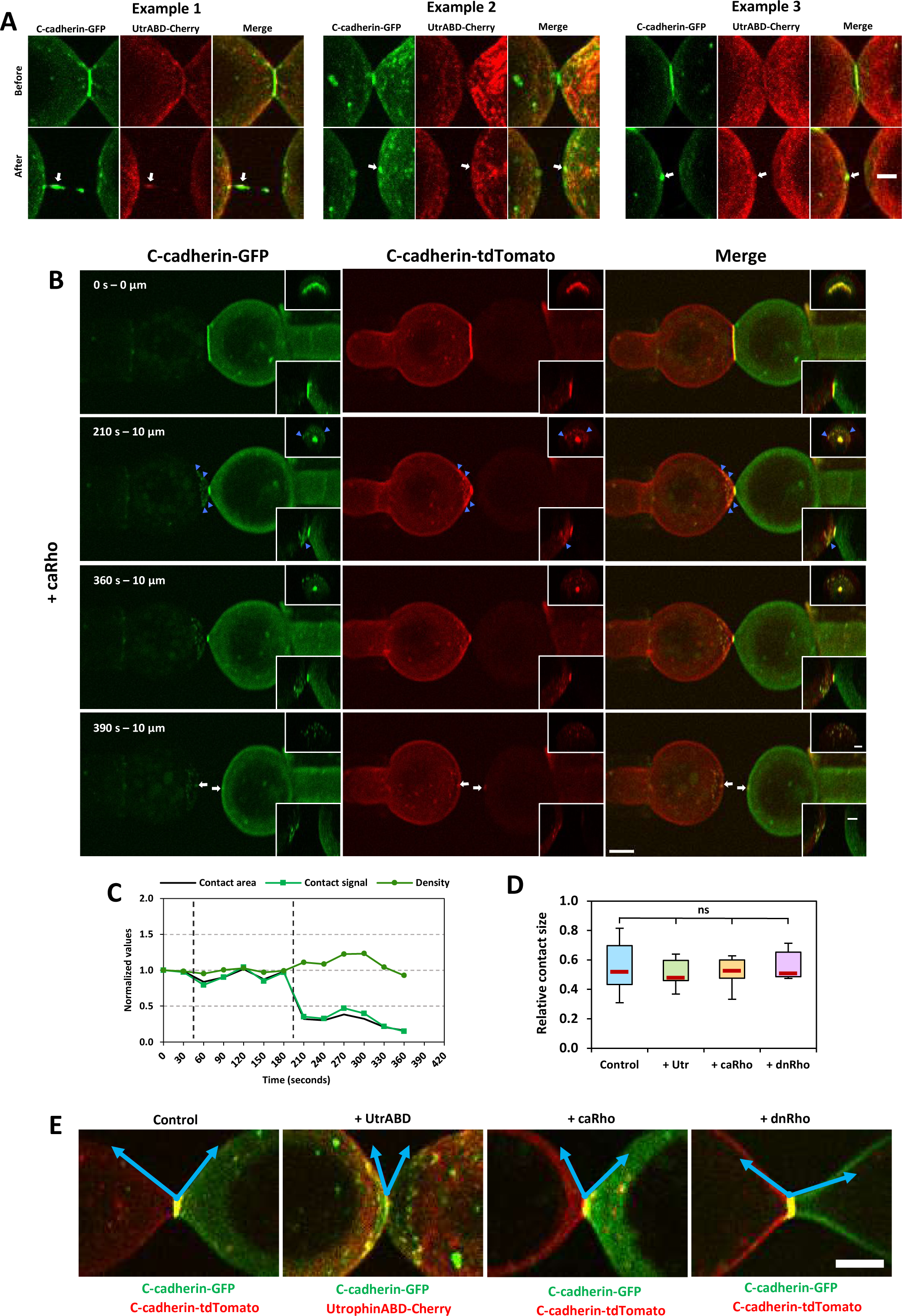
(related to Figure 6). A) Detailed view of final detachments of doublets of cell expressing UtrABD-Cherry, showing rupture of the remnant contact (C-cadherin-GFP, arrows). For each of the three examples, the frame just prior and just after detachment are shown. B) Example of massive early rupture for a doublet of two cells, both expressing caRho, and either C-cadherin-GFP or tdTomato, respectively. Blue arrowheads: Multiple clusters containing both green and red cadherin are observed at the surface of the left cell immediately after the second displacement (210 sec). The bright remnant of the contact is ruptured between 360 and 390 sec, without any further displacement. In this example, early rupture appears largely asymmetric, while late rupture is bilateral, leaving one spot on each cell membrane, each containing both green and red cadherin (white arrows). Top right insets are maximum intensity projections of the YZ orthogonal view. Low right insets are maximum intensity projections of the XZ orthogonal views. Scale bar for the main frame is 10 μm; insets are 5 μm. C) Quantification of contact and cadherin signal for the doublet showed in (B). Note the abrupt drop in contact surface and cadherin total signal at 210 sec (early rupture), followed by a small increase in cadherin density for the remaining shrinking contact. D) Comparison of the contact size before pipette pulling for the various conditions used in the DPA experiments. The relative size, expressed as contact diameter/cell diameter, appears similar in all conditions. (E) Examples of cell-cell contacts immediately prior to detachment for control, +UtrABD-Cherry, +caRho, and +dnRho doublets. For control, caRho, and dnRho cells on the right were expressing C-cadherin-tdTomato and cells on the left are expressing C-cadherin-GFP. UtrABD-Cherry cells were both expressing C-cadherin-GFP. White lines show the tangent along the membranes that were used to calculate the angle. Control and +dnRho have wide angles while + UtrABD and +caRho show more acute angles. Scale bar 10 μm.

**Supplemental Table S1.** List of mRNA used in this study with injected amounts

List of mRNA used in this study with injected amounts
| Plasmid | mRNA injected per blastomere at 2 cell stage (pg) |
| --- | --- |
| Constitutively active Alk4 (activin receptor type 1-B) | 1000 |
| Myc-β-catenin | 100 |
| C-cadherin-GFP | 600 |
| C-cadherin-dTomato | 600 |
| p120 catenin-GFP | 400 |
| α-catenin-GFP | 500 |
| Non-muscle myosin heavy chain 2A (NMHC2A)-YFP | 500 |
| Utrophin ABD-Cherry | 200 |
| Constitutively active RhoA (caRho) | 50 |
| Dominant negative RhoA (dnRho) | 150 |

All constructs were in the pCS2+ plasmid, except for Alk4, which was in pSP64T.

**Supplemental Table S2.**
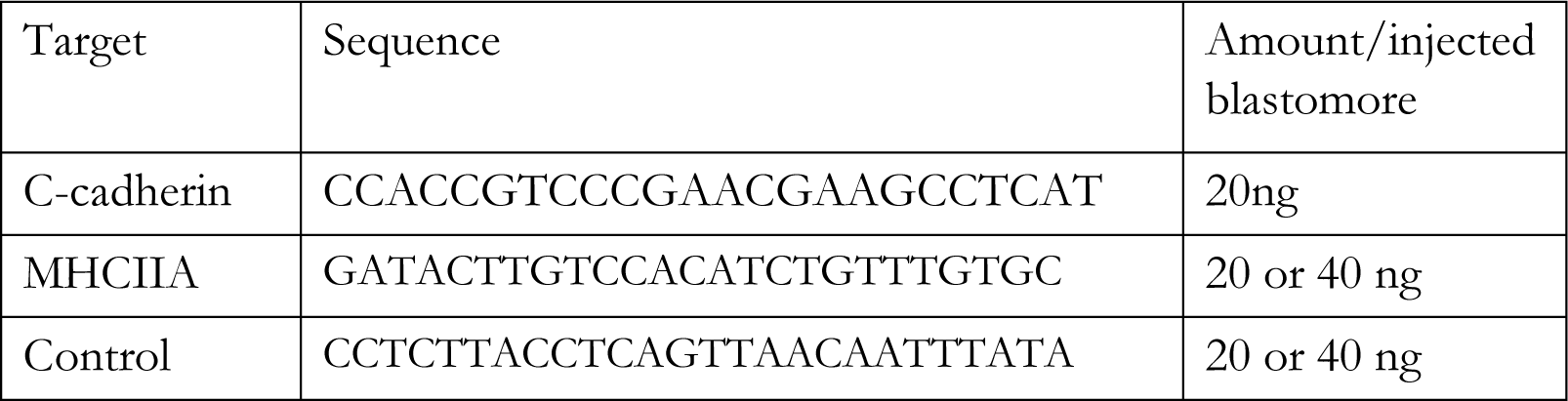
List of morpholinos with injected amounts

